# Transcriptional response to host chemical cues underpins expansion of host range in a fungal plant pathogen lineage

**DOI:** 10.1101/2020.10.29.360412

**Authors:** Stefan Kusch, Justine Larrouy, Heba M. M. Ibrahim, Noémie Gasset, Olivier Navaud, Malick Mbengue, Catherine Zanchetta, Celine Lopez-Roques, Cecile Donnadieu, Sylvain Raffaele

**Affiliations:** Laboratoire des Interactions Plantes-Microorganismes (LIPM), INRAE, CNRS, 24 Chemin de Borde Rouge - Auzeville, CS52627, F31326 Castanet Tolosan Cedex, France; Unit of Plant Molecular Cell Biology, Institute for Biology I, RWTH Aachen University, Worringerweg 1, D-52056 Aachen, Germany; Department of Pest Management and Conservation, Lincoln University, Lincoln 7647, Canterbury, New Zealand; Department of Genetics, Faculty of Agriculture, Cairo University, Gammaa St., 12613 Giza, Egypt; Division of Plant Biotechnics, KU Leuven, Willem de Croylan 42, BE-3001 Leuven, Belgium; INRAE, US 1426, GeT-PlaGe, Genotoul, Castanet-Tolosan, France, (doi:10.15454/1.5572370921303193E12)

**Author notes:** equal contribution (Justine Larrouy, Heba M. M. Ibrahim). **Correspondence:** Sylvain Raffaele.

**Keywords:** Mycotoxins, Plant defense, Regulatory variation, RNA sequencing, *Sclerotinia*, Virulence

## Abstract

The host range of parasites is an important factor in assessing the dynamics of disease epidemics. The evolution of pathogens to accommodate new hosts may lead to host range expansion, a process the molecular bases of which are largely enigmatic. The fungus *Sclerotinia sclerotiorum* parasitizes more than 400 plant species from diverse eudicot families while its close relative, *S. trifoliorum*, is restricted to plants from the *Fabaceae* family. We analyzed *S. sclerotiorum* global transcriptome reprogramming on hosts from six botanical families and reveal a flexible, host-specific transcriptional program driven by core and host-response co-expression (SPREx) gene clusters. We generated a chromosome-level genome assembly for *S. trifoliorum* and found near-complete gene space conservation in broad and narrow host range *Sclerotinia* species. However, *S. trifoliorum* showed increased sensitivity to the *Brassicaceae* defense compound camalexin. Inter-specific transcriptome analyses revealed a lack of transcriptional response to camalexin in *S. trifoliorum* and provide evidence that *cis*-regulatory variation associates with the genetic accommodation of *Brassicaceae* in the *Sclerotinia* host range. Our work demonstrates adaptive plasticity of a broad host range pathogen with specific responses to different host plants and demonstrates the co-existence of signatures for generalist and polyspecialist life styles in the genome of a plant pathogen. We reason that this mechanism enables the emergence of new disease with no or limited gene flow between strains and species, and could underlie the emergence of new epidemics originating from wild plants in agricultural settings.

## Introduction

The range of hosts that pathogens can infect is determined by genetic and environmental factors. This host range is an important factor in assessing the dynamics of disease epidemics. Specialists parasitize one or few hosts, such as the malaria parasite *Plasmodium falciparum* which is specific to humans (Otto et al., 2018) or the cereal powdery mildews infecting only cereals like rye and wheat (Troch et al., 2014). Generalists on the other hand have the ability to live on a range of diverse hosts, as for instance influenza A viruses infecting various Avian and Mammal species (Cauldwell et al., 2014). Natural selection by host populations and environmental factors drives frequent host switches and variations in pathogen host range. Specialization occurs when a pathogen adapts to a specific host and enters a co-evolutionary arms race with it. In some instances, adaptation following host switching and host jumps involves the ability to efficiently colonize new hosts while retaining the ability to colonize the original host lineage, resulting in host range expansions (Nylin and Janz, 2009; Thines, 2019).

Adaptation to the gene-for-gene type of plant resistance is a paradigmatic example of co-evolutionary arms race (Flor, 1971). Plant resistance genes typically recognize specific pathogenic proteins called effectors and mount a resistance reaction upon perception. Adapted pathogens evolved to avoid recognition by modification or loss of the respective effector (Jones and Dangl, 2006). This involves rapid adaptation, for example by selective sweeps (Sánchez-Vallet et al., 2018) leaving characteristic patterns of variation in the genome of plant pathogens (Möller and Stukenbrock, 2017; Raffaele and Kamoun, 2012). In the potato late blight pathogen *Phytophthora infestans*, the resulting genetic variation is notably responsible for a tradeoff in effector activity on targets from different hosts (Dong et al., 2014) and distinctive two-speed genome architecture (Dong et al., 2015; Raffaele et al., 2010). In addition, balancing selection occurs by genetic exchange between pathogen populations through gene flow and allows polymorphisms to persist in the gene pool and increases the genetic diversity in populations (Thrall et al., 2012). Arms-race models generally assume an isolated pathogen co-evolving with one host *via* pairwise selection. However, pathogen genomes often evolve in response to selection caused by more than one host under diffuse co-evolution (Ebert and Fields, 2020; Janzen, 1980). Genomic signatures of diffuse selection and molecular adaptations associated with interaction with multiple hosts are largely unresolved (Ebert and Fields, 2020).

Theories of evolutionary transitions suggest that genetic accommodation of pathogens to new hosts could entail general-purpose molecular bases supporting the colonization of any host (true generalist), or multiple modules dedicated to the colonization of specific hosts (polyspecialist) (Nylin and Janz, 2009; West-Eberhard, 1989). Comparative genomic studies have highlighted the role of expansion of gene families related to secondary metabolism and detoxification in host range expansion in insect (Simon et al., 2015) and fungal (Baroncelli et al., 2016) plant pathogens. A recent comparative study reported the transition from specialized one-speed genomes towards adaptive two-speed genomes correlated with increased host range in ergot fungi (Wyka et al., 2020). Similarly, analysis of seven fungi from the *Metarhizium* genus of entomopathogens showed an expansion in genes encoding G protein-coupled receptors, proteases, transporters, enzymes for detoxification and secondary metabolite biosynthesis, coinciding with increased host range (Hu et al., 2014). In this lineage, horizontal gene transfers contributed to host range expansion (Zhang et al., 2019a). By contrast, genome size was inversely correlated with host range in *Helicoverpa* butterflies (Zhang et al., 2019b). The genome of the generalist aphid *Myzus persicae* has a gene count half that of *Acyrthosyphon pisum*, which is specialized on pea. Instead, *M. persicae* colonizes diverse host plants through rapid transcriptional induction of specific gene clusters (Mathers et al., 2017). These studies support the polyspecialist model of host range expansion in which pathogen genomes harbor several specialized gene modules, resulting from gene family expansion or differential gene expression.

The white mold fungus *Sclerotinia sclerotiorum* is a typical generalist plant pathogen infecting more than 400 plant species (Boland and Hall, 1994). Its genome lacks signatures of selective sweeps (Derbyshire et al., 2019a) and two-speed architecture (Derbyshire et al., 2017). Host colonization by *S. sclerotiorum* is supported by division of labor enabling cooperation between cells of invasive hyphae (Peyraud et al., 2019). In addition, the *S. sclerotiorum* genome shows signatures of adaptive translation, a selective process shaping the optimization of codon usage to increase protein synthesis efficiency. Codon optimization is particularly clear in genes expressed during plant colonization and encoding predicted secreted proteins (Badet et al., 2017). Division of labor and codon optimization increase *S. sclerotiorum* fitness independently of the host genotype and therefore constitute genomic signatures of a true generalist. By contrast, a study of a few effector candidate genes suggested the existence of plant host-specific expression patterns (Guyon et al., 2014) and various small RNAs are differentially expressed on *A. thaliana* and common bean *(Phaseolus vulgaris)* (Derbyshire et al., 2019b). Furthermore, various isoforms originating from host-specific alternative splicing add additional plasticity to the transcriptomic profile of *S. sclerotiorum* (Ibrahim et al., 2020) and suggest polyspecialist adaptations in this species. Nevertheless, the extent to which the *S. sclerotiorum* genome harbors signatures of adaptation to a polyspecialist lifestyle has not been fully elucidated.

To test whether *S. sclerotiorum* activates distinct gene sets for the infection of diverse host species, we took advantage of transcriptomic expression data obtained from *in vitro* cultures of *S. sclerotiorum* and during infection of plants covering major botanical families in the Pentapetalae (Ibrahim et al., 2020; Peyraud et al., 2019; Sucher et al., 2020). We reveal a flexible, host-specific transcriptional program driven by core and host-response co-expression (SPREx) clusters. Genome sequencing and assembly revealed that the sister species *S. trifoliorum*, unable to infect *Brassicaceae* plants, exhibits high level of gene space conservation. We show that lack of transcriptional response to camalexin coincides with increased sensitivity of *S. trifoliorum* to this *Brassicacea* defense compound. Interspecific comparison of promoter sequences provide evidence that *cis*-regulatory variation associates with the genetic accommodation of *Brassicaceae* in the *Sclerotinia* host range. Our work demonstrates adaptive plasticity of a broad host range pathogen with specific responses to different host plants and exemplifies the co-existence of signatures for generalism and polyspecialism in the genome of a plant pathogen.

## Results

### A subset of host species trigger specialized transcriptome reprogramming in *S. sclerotiorum*

To investigate transcriptional reprogramming in *S. sclerotiorum* during the colonization of hosts from different botanical families, we performed RNA-seq analysis of *S. sclerotiorum* 1980 during infection of thale cress *(Arabidopsis thaliana, Ath)*, tomato *(Solanum lycopersium, Sly)*, sunflower *(Helianthus annuus, Han)*, common bean *(Phaseolus vulgaris, Pvu)*, castor bean *(Ricinus communis, Rco)*, and beetroot *(Beta vulgaris, Bvu).* To control for variations in the kinetics of pathogen colonization on different hosts, samples were collected at similar infection stages by macro-dissection of disease lesion center and edge (Peyraud et al., 2019) in biological triplicates (36 samples in total). Percentage of reads mapped to *S. sclerotiorum* was 64.8% ± 4.0 in edge samples, 90.4% ± 2.2 in center samples (**Figure 1A**). We updated the *S. sclerotiorum* 1980 gene annotation and identified 360 genes as likely transposable elements with RepeatMasker (Smit et al., 2016) that were excluded from further analysis, resulting in 10,770 protein-coding genes in *S. sclerotiorum* (**Supplementary Table 1)**. We filtered out genes weakly or not expressed by excluding genes with FPKM < 25 in all 36 plant infection samples and in *S. sclerotiorum* mycelium grown *in vitro* in PDB and on PDA plates, leaving 7,524 genes for differential expression (DE) analysis **(Supplementary Table 2)**. The center and edge of mycelium grown in PDA were used as references in DE analysis for center and edge of lesion, respectively, leading to the identification of 2,625 DEGs in total. We identified 1,120 and 1,209 genes significantly up-regulated (log_2_ fold change LFC ≥ 1; adjusted p-value ≤ 0.01) in lesion edge and center, respectively (**Figure 1B**, **Supplementary Table 3**) across six hosts. There was a total of 592 and 1,103 genes significantly down-regulated in lesion edge and center respectively across six hosts.

**Figure 1.**
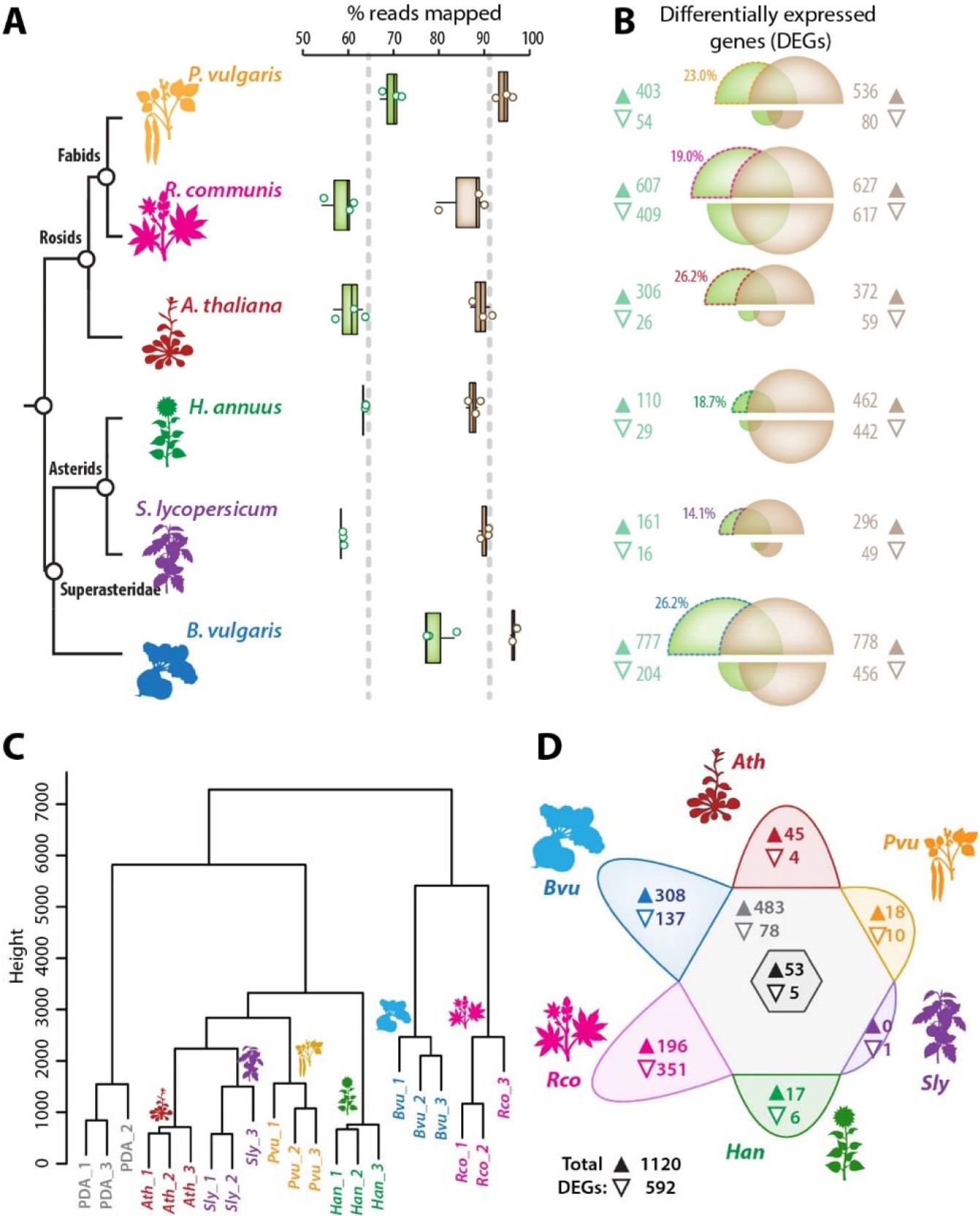
Differential regulation of *S. sclerotiorum* transcriptome during the colonization of plants from six botanical families. **(A)** Proportion of RNA sequencing reads from colony edge samples (green) and colony center samples (brown) mapped to a unique position in the *S. sclerotiorum* genome (% of all reads). Results from samples collected in three independent biological replicates are shown, grey dashed line shows average for all samples. Boxplots show 1st and 3rd quartiles (box), median (thick line) and the most dispersed values within 1.5 times the interquartile range (whiskers). **(B)** Number of *S. sclerotiorum* genes expressed differentially (DEGs) on each host compared to *in vitro-grown* colonies. Bubbles of the venn diagrams are sized proportionally to the number of DEGs in each treatment, labels on the right and left side showing the corresponding number of genes. DEGs detected in colony edge are shown in green, those in colony center are shown in brown. The upper half of the bubbles corresponds to genes upregulated, the lower half to genes downregulated. The proportion of upregulated genes unique to edge samples is labelled on diagrams and the corresponding sectors highlighted by dotted lines. **(C)** Hierarchical clustering of edge samples based on the expression of 2,625 DEGs. Numbers in branch labels correspond to biological replicates. **(D)** Distribution of DEGs according to host species. For each sector, the upper value (▲) correspond to upregulated genes, the lower value (▽) to downregulated genes. The central dark grey hexagon shows DEGs detected on all six host, the light grey hexagon shows DEGs detected on 2 to 5 hosts. *Ath, Arabidopsis thaliana; Bvu, Beta vulgaris; Han, Helianthus annuus; Pvu, Phaseolus vulgaris; Rco, Ricinus communis; Sly, Solanum lycopersicum.*

We noted a strong variation in the number of *S. sclerotiorum* DEGs on the various host plants (**Figure 1B, Supplementary Table 4**). The number of upregulated genes in lesion edge ranged from 110 on *H. annuus* to 777 on *B. vulgaris* (7.1-fold variation) and the number of down-regulated genes in lesion edge ranged from 16 on *S. lycopersicum* to 409 on *R. communis* (25.6 fold variation). The number of *S. sclerotiorum* DEGs in lesion center varied between hosts to a lesser extend (2.6 and 12.6 fold variation for genes up- and down-regulated, respectively). As suggested by (Peyraud et al., 2019), the degree of differentiation between edge and center cells varied according to the host, with 14.1% of DEGs specific to fungal edge cells on *S. lycopersicum* and up to 26.2% on *A. thaliana* and *B. vulgaris.* Hierarchical clustering (**Figure 1C**) and principal component analysis (**Supplementary Figure 1**) using FPKM values for the 2,625 DEGs showed clear separation of *S. sclerotiorum* gene expression in edge samples according to the infected host (**Supplementary Table 5**). In colony edge, 53 DEGs (4.7%) were upregulated during the colonization of all host plants, 483 DEGs (43.1%) where upregulated on at least two host plants, and 584 DEGs (52.1%) showed specific upregulation on one host (**Figure 1D**). Similarly, 5 DEGs (0.8%) were downregulated during the colonization of all host plants, 78 DEGs (13.2%) were downregulated on at least two host plants, and 509 DEGs (86%) were downregulated during the colonization of one host only. The number of genes upregulated on one host only represented 0% on *S. lycopersicum*, 4.5% on *P. vulgaris*, 14.7% on *A. thaliana*, 15.5% on *H. annuus*, 32.3% on *R. communis* and 39.6% on *B. vulgaris.* These results suggest that the colonization of *S. lycopersicum* relies mostly on *S. sclerotiorum* core virulence genes while the colonization of *A. thaliana, R. communis* and *B. vulgaris* may require the activation of host-specific fungal virulence programs.

### The functional landscape of *S. sclerotiorum* core and host-specific transcriptome

To document the functional diversity of *S. sclerotiorum* genes differentially expressed during host colonization, we analyzed GO and PFAM annotation enrichment with genes upregulated *in planta.* We detected 229 GO terms in the 7,423 expressed genes, distributed among 6,164 genes, leaving 1,259 genes (16.9%) with no GO annotation. In addition to the three root ontologies, used for the annotation of genes with unknown function, we found 11 GOs significantly enriched with upregulated genes (chi-squared adjusted p-val < 0.01) during the colonization of at least one plant (**Supplementary Table 6**). The six ontologies enriched in genes upregulated during the colonization of multiple host species related to the metabolism of cell wall compounds (GO:0016798, GO:0005975, GO:0044723), redox homeostasis (GO:0016491) and the extracellular space (GO:0005615, GO:0005576). The ontology “pathogenesis” (GO:0009405) was enriched in genes upregulated during infection of tomato, “secondary metabolic process” (GO:0019748) and three ontologies related to membrane transport (GO:0016020, GO:0016021, GO:0022857) were enriched in genes upregulated during infection of beetroot.

We detected 8,764 PFAM domains in the 7,423 expressed genes, distributed among 5,746 genes, leaving 1,677 genes (22.6%) with no PFAM annotation. We found 94 PFAMs significantly enriched with upregulated genes (chi-squared adjusted p-val < 0.01) during the colonization of at least one plant (**Figure 2A, Supplementary Table 7**). Thirteen of these PFAMs were enriched in genes upregulated during the colonization of five or six host species, including galactosidase (PF10435, PF13363, PF13364, PF16499), glycosyl hydrolase (PF00150, PF01301, PF00295), sugar transporter (PF00083), cytochrome P450 (PF00067) and domain of unknown function DUF4965 (PF16335) domains. Forty PFAMs were enriched in genes upregulated during the colonization of a single host species. This included glycosyl hydrolase domains (PF00723 and PF00232 on *A. thaliana*, PF07745 on *R. communis*, PF03443 and PF00457 on *B. vulgaris).* Several domains were likely related to detoxification processes, such as the condensation domain (PF00668) enriched on *P. vulgaris* and involved in non-ribosomal peptide biosynthesis, the stress responsive A/B barrel domain (PF07876) and the HpcH/HpaI aldolase/citrate lyase family domain (PF03328) enriched on *R. communis* and the Epoxide hydrolase domain (PF06441) enriched on *H. annuus.*

**Figure 2.**
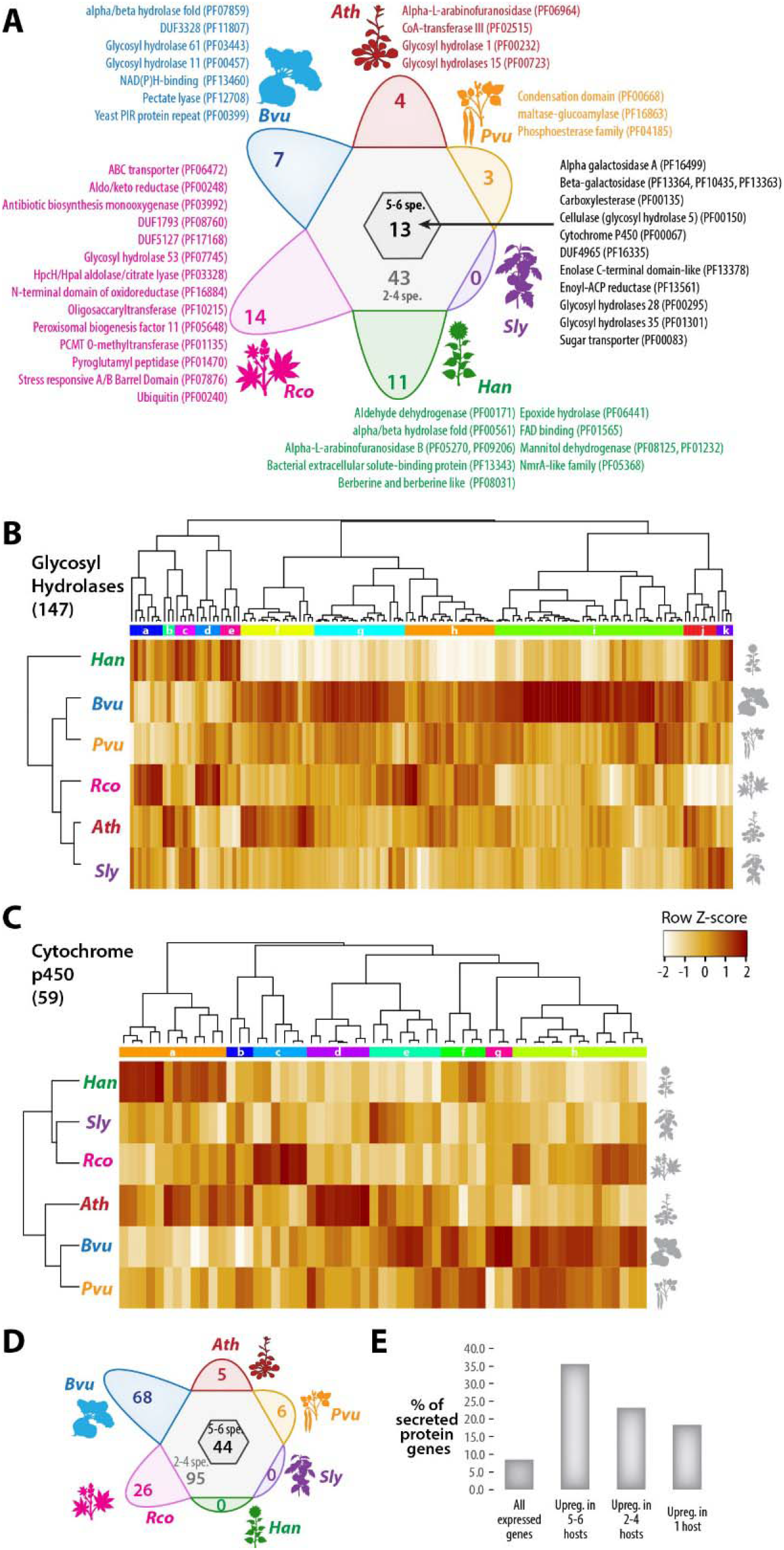
Distribution of *S. sclerotiorum* functional gene groups according to hosts in which they are transcriptionally upregulated. **(A)** PFAM domains enriched with upregulated genes according to host. Domains enriched with genes upregulated on one host only or five and six hosts are labelled on the figure, with font color corresponding to host species. **(B)** Hierarchical clustering of log_2_ fold change of expression for 147 expressed genes encoding glycosyl hydrolases. Eleven hierarchical clusters labelled a-k were delimited. **(C)** Hierarchical clustering of log_2_ fold change of expression for 59 expressed genes encoding cytochrome p450. Eight hierarchical clusters labelled a-h were delimited. **(D)** Distribution of genes encoding predicted secreted proteins according to host species in which they are upregulated. **(E)** Proportion of genes encoding predicted secreted proteins among all expressed genes and genes upregulated *in planta. Ath, Arabidopsis thaliana; Bvu, Beta vulgaris; Han, Helianthus annuus; Pvu, Phaseolus vulgaris; Rco, Ricinus communis; Sly, Solanum lycopersicum.*

Next, we examined the differential expression pattern of the largest gene families enriched in upregulated genes using hierarchical clustering of LFC values. We identified 147 expressed genes harboring a glycosyl hydrolase (GH) domain that classified into 11 hierarchical clusters (a to k, **Figure 2B**). The colonization of each host upregulated specific sets of GH genes. For instance, the colonization of *H. annuus* activated mostly GH genes from clusters a, b, c, d, e, j and k. The colonization of *B. vulgaris* activated mostly GH genes from clusters f, g, h and i, while the colonization of *A. thaliana* activated mostly GH genes from clusters b, c, f and j. We identified 59 expressed genes harboring a cytochrome p450 domain (Peyraud et al., 2019) into 8 hierarchical clusters (a to h, **Figure 2C**). Here again, the colonization of each host lead to a specific signature of p450 genes upregulated in *S. sclerotiorum.* Genes upregulated belonged to clusters a and f on *H. annuus*, e on *S. lycopersicum*, c on *R. communis*, a, b and d on *A. thaliana.* The 1,120 upregulated genes included 246 genes (21.9%) coding for putative secreted proteins, among which 44 (17.9%) were upregulated on five or six hosts, 95 (38.6%) were upregulated on two, three or four hosts, and 105 (42.6%) were upregulated on one host only (**Figure 2D**). Genes encoding putative secreted proteins represented 8.5% of the 7,423 expressed genes, 18.3% of genes upregulated on one host, 23.2% of genes upregulated on two, three or four hosts and 35.5% of genes upregulated on five or six hosts (**Figure 2E**). This enrichment is consistent with a prominent role of fungal secreted proteins in the interaction with host plants and indicates that secreted proteins contribute to a larger part of *S. sclerotiorum* core infection program than host-specific infection programs.

### *S. sclerotiorum* transcriptome is organized in generalist and host-specialized genomic clusters

Several fungal genomes harbor biosynthetic clusters containing co-expressed genes (Ámon et al., 2017; Wisecaver and Rokas, 2015). To determine the extent to which *S. sclerotiorum* genes expressed during plant infection are organized in genomic clusters, we analyzed local gene expression correlation during the infection of six host plants using a sliding window approach (**Supplementary Table 8**). Average Pearson correlation of gene expression within 9-gene windows was 0.077 ± 0.076 at the whole genome scale. 2,222 windows had Pearson correlation ≥0.15, 634 of which (28.5%) residing in 58 loci that we designated as *Sclerotinia* plant-response co-expression (SPREx) clusters distributed along the 16 chromosomes of *S. sclerotiorum* (**Figure 3A**). SPREx clusters included 5 to 46 genes, with an average local expression correlation between 0.136 and 0.298 (**Supplementary Table 9**). Twenty-five SPREx clusters (43.1%) overlapped at least partly with predicted secondary metabolite biosynthetic clusters identified with AntiSMASH (Graham-Taylor et al., 2020), 48 SPREx clusters (82.8%) included at least one gene significantly induced *in planta.*

**Figure 3.**
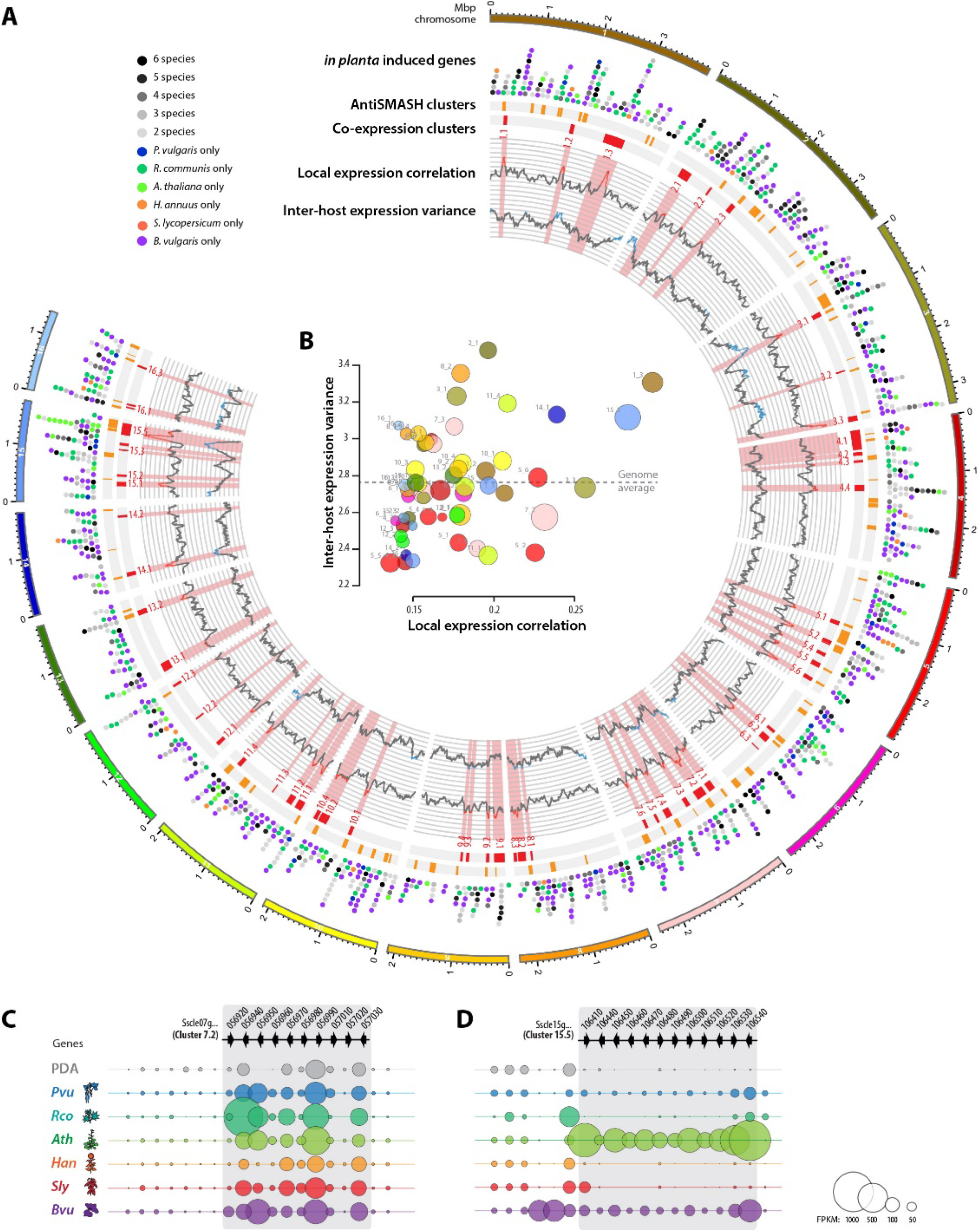
Genomic architecture of *S. sclerotiorum* infection transcriptome. **(A)** Distribution of gene expression information along the 16 chromosomes of *S. sclerotiorum.* From outermost to innermost track: *S. sclerotiorum* chromosomes with sequence length indicated in million base pairs (Mbp); position of *in planta* induced genes represented by circles colored according to hosts in which the gene is induced; position of secondary metabolite biosynthetic clusters identified by AntiSMASH; position of *Sclerotinia* plant response co-expression (SPREx) clusters identified in this work; local expression correlation along chromosomes, highlighted in red in regions where ≥0.15; inter-host expression variance along chromosomes (log scale), highlighted in blue in regions where ≥2.739. **(B)** Distribution of SPREx clusters according to their average local expression correlation (X-axis) and inter-host expression variance (log scale, Y-axis). Bubbles are sized according to the number of genes in SPREx clusters, from 5 to 43, and colored according to the chromosome they belong to (same color code as in A). **(C)** Expression of a subset of genes from the core SPREx cluster 7.2. The position and orientation of genes is indicated by arrowheads, labelled with gene names. The size of bubbles shows the average FPKM values for each gene in six conditions. **(D)** Same as C for the host-specific SPREx cluster 15.5. *Ath, Arabidopsis thaliana; Bvu, Beta vulgaris; Han, Helianthus annuus; Pvu, Phaseolus vulgaris; Rco, Ricinus communis; Sly, Solanum lycopersicum.*

To gain an overview of the consistency of *S. sclerotiorum* gene expression across hosts, we calculated the inter-host expression variance for each gene using FPKM values in edge samples during the infection of six host plants (**Supplementary Table 8-9**). Average log of inter-host expression variance was 2.739 at the genome level and ranged from 2.325 to 3.481 for the 58 SPREx clusters (**Figure 3B**). This identified SPREx clusters in which genes are expressed stably in all colonized hosts (inter-host expression variance lower than genome average), such as cluster 7.2. This expression cluster includes a secondary metabolite biosynthetic cluster (Sscle07g056450 to Sscle07g056660) weakly expressed *in planta*, and a region with genes induced in planta and genes encoding predicted secreted proteins (Sscle07g056950 - Sscle07g056990, Sscle07g057030) (**Figure 3C**). We also identified SPREx clusters harboring genes with expression highly variable according to host (inter-host expression variance higher than genome average), such as cluster 15.5. This SPREx cluster overlaps with two secondary metabolite biosynthetic clusters (Sscle15g106410 to Sscle15g106590, orthologous to *B. cinerea* botcinic acid biosynthetic cluster (Dalmais et al., 2011; Graham-Taylor et al., 2020; Peyraud et al., 2019), and Sscle15g106630 to Sscle15g107150) and includes seven genes encoding predicted secreted proteins. Genes of SPREx cluster 15.5 corresponding to botcinic acid biosynthesis cluster are highly expressed during the colonization of *A. thaliana* but not the other five plant species we tested (**Figure 3D**).

To gain insights into biological processes associated with the physical clustering of genes in the *S. sclerotiorum* genome, we analyzed gene ontologies and PFAM domains enriched with genes present in SPREx clusters (**Supplementary Table 10**). Eighty-one GOs were significantly enriched with genes in SPREx clusters (chi-squared test adjusted p-val < 0.01), including 30 “biological process” GOs, 20 “cellular component” GOs and 31 “molecular function” GOs. Eleven PFAM domains were significantly enriched with genes in SPREx clusters. The enriched annotations were notably related to secondary metabolism (GO:0019748 “secondary metabolic process”, PF00109, PF16197 and PF02801 “Ketoacyl synthase”, PF14765 “Polyketide synthase dehydratase”, PF00698 “Acyl transferase”), cytoskeleton dynamics (GO:0008092 “cytoskeletal protein binding”, GO:0005856 “cytoskeleton”, PF00307 “Calponin homology”), cell wall remodeling (GO:0005975 “carbohydrate metabolic process”, GO:0016798 “hydrolase activity, acting on glycosyl bonds” PF00187 “Chitin recognition protein”, GO:0005618 “cell wall”), and cellular signaling (PF00071 “Ras family”, GO0006950 “response to stress”, GO0007165 “signal transduction”, GO0001071 “DNA-binding transcription factor activity”).

### *S. trifoliorum* SwB9 genome assembly and annotation reveals a nearly complete conservation of *S. sclerotiorum* genes induced *in planta*

*S. trifoliorum* is closely related to *S. sclerotiorum* but has a host range restricted to plants in the Asterales and Fabids families (Navaud et al., 2018). Unlike *S. sclerotiorum, S. trifoliorum* colonizes *A. thaliana* very poorly to not at all. To gain insights into the evolution of *A. thaliana-specific* upregulated genes in the *Sclerotinia* lineage, we performed genome sequencing of *S. trifoliorum* isolate SwB9 using Illumina short-read and nanopore long-read data. The final genome assembly of *S. trifoliorum* SwB9 comprised 42 contigs totaling 39,909,921 bp. The assembly contained 32 cases of canonical telomeric repeat sequences (5’-TTAGGG-3’ hexamer; 15 to 28 tandem copies), suggesting that like *S. sclerotiorum S. trifoliorum* SwB9 possesses 16 chromosomes. We assessed completeness of the genome with BUSCO (Simão et al., 2015) and found 97.4% of highly conserved ascomycete genes present, suggesting near-complete gene space coverage. According to RepeatMasker v4.0.7, around 3.7% of the genome are repetitive or made up of transposable elements, which is somewhat lower than the 6.2% found in *S. sclerotiorum* (Derbyshire et al., 2017). Three contigs (tig00000348, tig00000347, and tig00000297) had BLASTN hits for mitochondrial DNA, indicating that these contigs belong to the *S. trifoliorum* SwB9 mitochondrion with a total length of at least 226,016 bp.

According to synteny analysis with MUMmer3 (Kurtz et al., 2004), the genomes share a high level of collinearity (**Figure 4A**), and 67.2% of the *S. trifoliorum* genome aligned to 68.8% of the *S. sclerotiorum* genome. While most *S. sclerotiorum* chromosomes are syntenic, we found cases of inversions as well as an apparent chromosome arm exchange of chromosomes 6 and 12 in the genome of *S. trifoliorum* on the corresponding contigs tig00000041 and tig00005977 (**Figure 4B**). Overall, we generated a near-chromosome assembly for *S. trifoliorum* SwB9 of sufficient quality for gene space comparison with *S. sclerotiorum.*

**Figure 4.**
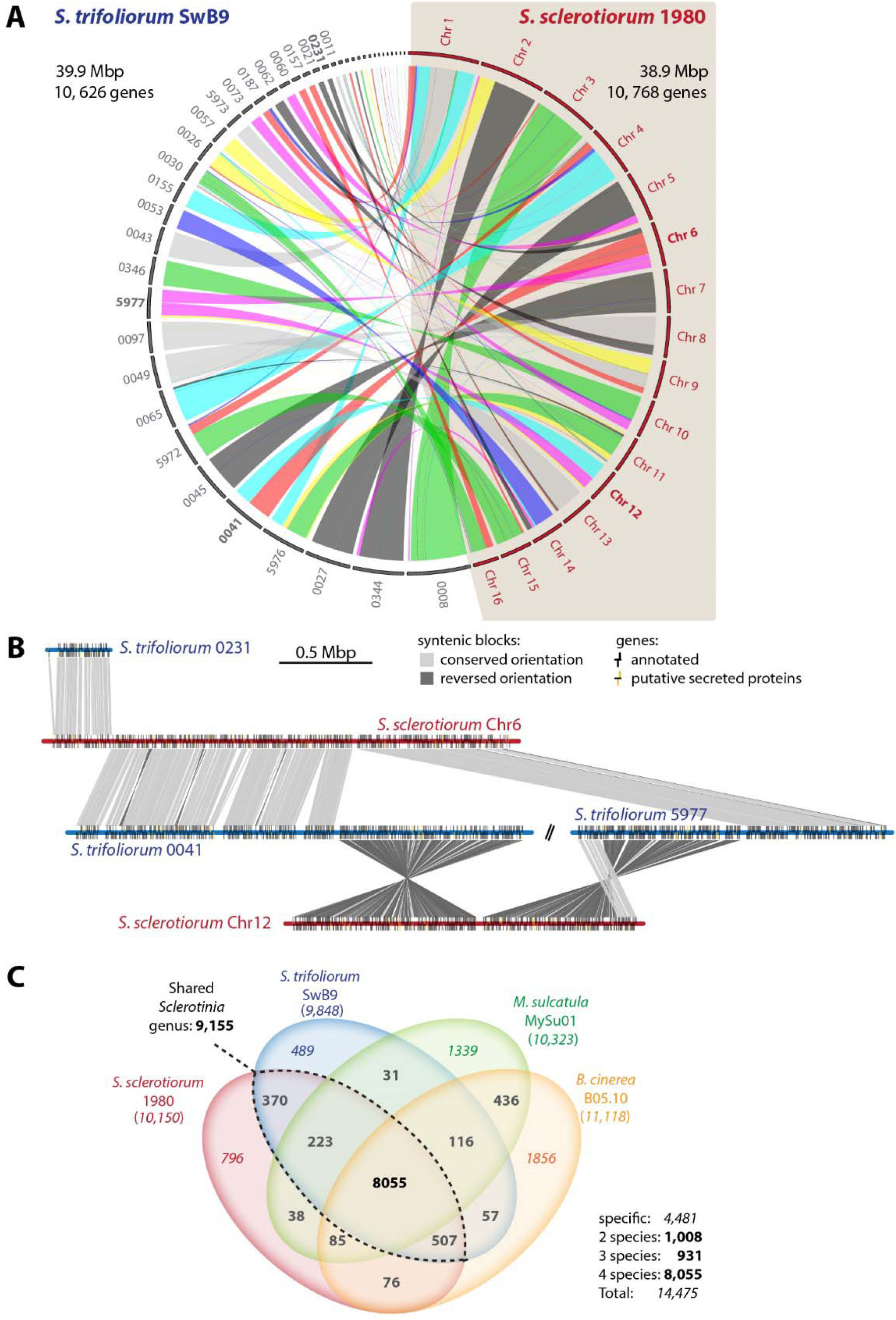
The genome of *S. trifoliorum* resembles the genome of *S. sclerotiorum.* We generated a near-chromosome level assembly for *S. trifoliorum* SwB9. (**A**) The synteny of the two genomes was analyzed using MUMmer (Kurtz et al., 2004). The circos plot shows the synteny between the chromosomes of *S. sclerotiorum* 1980 and the newly assembled contigs of *S. trifoliorum* SwB9. Chromosomes and contigs shown in B are labelled with bold fonts. (**B**) Synteny of *S. sclerotiorum* chromosomes 6 and 12 against the *S. trifoliorum* assembled contigs plotted by genoPlotR (Guy et al., 2010). (**C**) Venn diagram summarizing the results from the Orthofinder analysis between the proteomes of *S. sclerotiorum* 1980, *S. trifoliorum* SwB9, *Myriosclerotinia sulcatula* MySu01, and *Botrytis cinerea* B05.10. Values in bold correspond to number of orthogroups, values in italics correspond to number of genes.

Next, we used BRAKER2 (Hoff et al., 2016) for unsupervised *ab initio* gene prediction based on RNA-seq. BRAKER2 predicted 11,290 gene models (11,283 excluding removed and mitochondrial contigs). We then used WebApollo (Lee et al., 2013) for manual curation, resulting in 10,626 unique gene models in the *S. trifoliorum* SwB9 annotation, 28 of which are incomplete (lacking start or stop codon) due to terminal location in their contig. The *S. trifoliorum* proteome comprises 2,020 proteins with putative transmembrane domains, and according to PFAM prediction using hmmscan (**Supplementary Table 11**), the most abundant domains are transporter domains (MFS1 and AAA-type), kinases, alcohol dehydrogenase, and Zn-cluster. 705 proteins contained predicted secretion signal peptides, 73 of which are possible effector candidates according to an EffectorP 2.0 search (Sperschneider et al., 2018).

Using OrthoFinder (Emms and Kelly, 2015) and including *B. cinerea* B05.10 (Van Kan et al., 2017) and *Myriosclerotinia sulcatula* MySu01 (Kusch et al., 2020) annotated proteomes in the analysis, we found 9,155 orthogroups (OGs) shared between the two *Sclerotinia* species containing 10,085 (*S. sclerotiorum)* and 9,922 (*S. trifoliorum)* genes, respectively (**Figure 4C** and **Supplementary Table 12**). 8,055 OGs were common for all four species, 408 of which (5.3%) contained putative secreted proteins. Altogether, the genomes of the broad host spectrum fungus *S. sclerotiorum* and the narrow host spectrum fungus *S. trifoliorum* are highly syntenic and harbor a conserved proteome encoded by more than 9,900 core genes. 685 *S. sclerotiorum* genes had no ortholog in *S. trifoliorum*, including 249 *S. sclerotiorum* genes from 199 OGs that did not contain *S. trifoliorum* genes. Out of 306 *S. sclerotiorum* genes induced at the edge of colonies on *A. thaliana*, 300 (98%) had orthologs in *S. trifoliorum.* Out of 45 *S. sclerotiorum* genes upregulated specifically on *A. thaliana*, 42 (93.3%) had orthologs in *S. trifoliorum.* Therefore, expansion of the host range of Sclerotiniaceae fungi to *A. thaliana* largely relied on genes acquired prior to the divergence between *S. sclerotiorum* and *S. trifoliorum.*

### *S. trifoliorum* is highly sensitive to phytoalexins

Plants in the order Brassicales like *A. thaliana* produce tryptophan-derived defensive metabolites such as indolic glucosinolates (iGLs) and the indole alkaloid camalexin (cam.). The ability of some insect and fungal pathogens to metabolize these plant defense compounds into non-toxic derivatives contributes to their capacity to infect plants from the Brassicales (Chen et al., 2020). To test whether the toxicity of tryptophan-derived plant defense metabolites could explain the inability of *S. trifoliorum* to colonize Brassicales, we compared the sensitivity of *S. sclerotiorum* and *S. trifoliorum* to the five major defense tryptophan-derivatives produced by *A. thaliana* using an *in vitro* growth assay (**Figure 5A, Supplementary Figure 2**). Overall, radial growth of *S. trifoliorum* on PDA was slower than that of *S. sclerotiorum.* Nevertheless, both *S. sclerotiorum* and *S. trifoliorum* tolerated similar concentrations of tryptophan, raphanusamic acid, indole-3-carboxylic acid and indole-3ylmethylamine. However, the growth of *S. trifoliorum* was completely inhibited by 125 μM camalexin and 250 μM brassinin, while *S. sclerotiorum* remained able to grow. To verify that these compounds contribute to plant resistance to *S. trifoliorum*, we compared the colonization of wild type and *cyp79b2/b3 A. thaliana* plants, defective in iGLs biosynthesis (Stotz et al., 2011) (**Figure 5B**). Three days post inoculation (dpi), *S. sclerotiorum* had fully colonized 56% of Col-0 wild type leaves, while *S. trifoliorum* hardly grew out of the inoculation plug (0% of leaves fully colonized), consistent with the inability of *S. trifoliorum* to colonize *A. thaliana.* The *cyp79b2/b3* mutant was more susceptible to *S. sclerotiorum*, harboring 89% of leaves fully colonized at 3 dpi. Remarkably, *cyp79b2/b3* plants appeared susceptible to *S. trifoliorum* in this assay, as 67% of leaves were fully colonized at 3 dpi. These results indicate that sensitivity to tryptophan-derived defense metabolites contribute to the inability of *S. trifoliorum* to infect *A. thaliana.*

**Figure 5.**
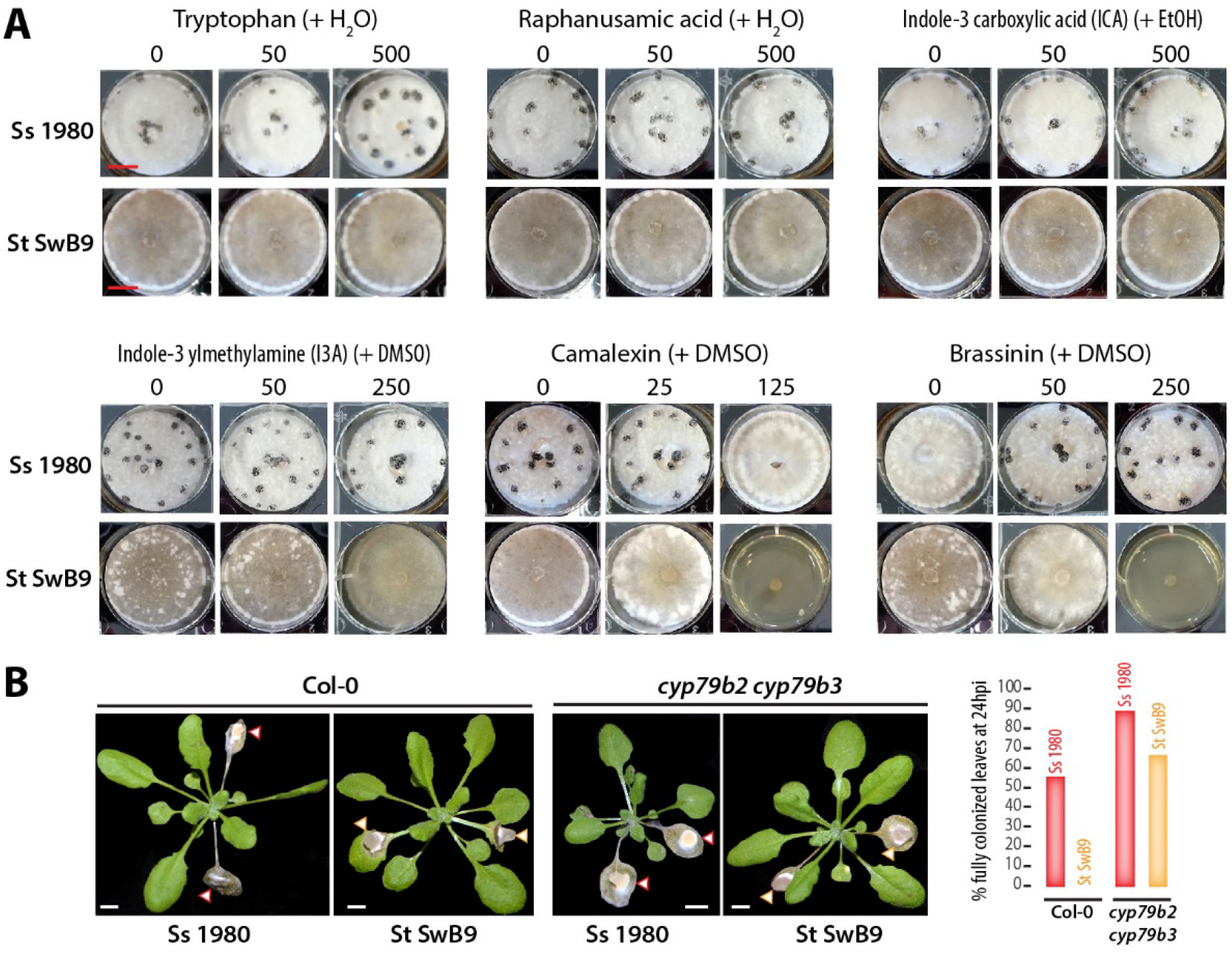
*S. trifoliorum* is more sensitive than *S. sclerotiorum* to phytoalexins *in vitro* and *in planta.* (**A**) Phytoalexin tolerance plate assay. *S. sclerotiorum* 1980 and *S. trifoliorum* SwB9 were cultivated on potato dextrose agar (PDA) containing phytoalexins at different concentrations. The solvent used for each compound is indicated between brackets. Additional concentrations and solvent-only conditions are shown in **Supplementary Figure 2**. Photos were taken after seven days; the experiment was conducted three times with similar results. Scale bar: 1 cm. (**B**) The *A. thaliana* wild type Col-0 and the indole glucosinolate and camalexin deficient mutant *cyp79b2 cyp79b3* were inoculated with *S. sclerotiorum* 1980 and *S. trifoliorum* SwB9, respectively. Photos were taken three days after inoculation. Arrowheads indicate agar plugs with *S. sclerotiorum* (red) or *S. trifoliorum* (yellow). Scale bar: 1 cm. The bar chart indicates the proportion of inoculated leaves fully colonized by each fungus for n=9 or 10 leaves.

### *S. sclerotiorum* but not *S. trifoliorum* reprograms its transcriptome in response to camalexin

In spite of the very high synteny of *S. sclerotiorum* and *S. trifoliorum* genomes, we observed a significant difference in sensitivity to camalexin and brassinin between these two species. *S. sclerotiorum* uses the isothiocyanate hydroxylase *SsSaxA (Sscle05g040340)* to detoxify plant defense compounds derived from glucosinolates (Chen et al., 2020). *S. sclerotiorum* also uses the brassinin glucosyltransferase *SsBGT1 (Sscle01g003110)* to metabolize brassinin (Pedras et al., 2004; Sexton et al., 2009).

These two genes are strongly induced during *A. thaliana* colonization, and both possess clear orthologs in *S. trifoliorum* genome. None of *S. sclerotiorum* genes induced during the colonization of *A. thaliana* and absent from *S. trifoliorum* genome could be unambiguously associated with detoxification of phytoalexins, suggesting that differences in their ability to colonize plants from the Brassicales is not related to differences in the enzyme-coding complement of their genome. *SsBGT1* was identified thanks to its strong induction in response to phytoalexins (Sexton et al., 2009). This prompted us to explore the extent to which *S. sclerotiorum* and *S. trifoliorum* differ in their transcriptional response to camalexin. For this, we performed RNA sequencing of *S. sclerotiorum* and *S. trifoliorum* colonies grown on PDA plates with camalexin. We used camalexin at 125 μM and 25 μM for *S. sclerotiorum* and *S. trifoliorum*, respectively, corresponding to the highest concentration of camalexin that supported the growth of each species in our assays, and colonies grown on PDA with DMSO as controls. The number of reads uniquely mapped to the *S. sclerotiorum* reference genome ranged from 9,658,323 to 21,158,861 (**Supplementary Table 13**). Out of the 7,524 *S. sclerotiorum* expressed genes we identified previously, 6,659 were expressed with FPKM≥25 in at least one sample of this assay. We performed DE analysis using limma on these 6,659 *S. sclerotiorum* expressed genes. At LFC≥1 and adjusted p-value≤0.01, 323 genes were up-regulated in *S. sclerotiorum* on camalexin (**Supplementary Table 14**). Among those, 180 (55.7%) were also induced at the edge of colonies in at least one of the plant species (**Supplementary Table 14**) and 301 had orthologs in *S. trifoliorum.* The number of reads uniquely mapped to the *S. trifoliorum* reference genome ranged from 16,356,783 to 22,673,171 (**Supplementary Table 13**). We detected 4,885 *S. trifoliorum* genes expressed at FPKM≥25 in at least one sample of this assay and performed DE analysis using limma on these genes. At LFC≥1 and adjusted p-val∪tó¤.01, 42 genes were up-regulated in *S. trifoliorum* on camalexin (**Supplementary Table 14**), among which 40 had orthologs in *S. sclerotiorum.* Hierarchical clustering of expression of the 341 DEGs and their orthologs from the two species identified five major clusters (a to e) **(Figure 6A**). Cluster a included genes upregulated by camalexin in *S. trifoliorum* only, cluster c included few genes repressed by DMSO, cluster d included genes upregulated by camalexin both in *S. sclerotiorum* and in *S. trifoliorum*, cluster e included genes upregulated by camalexin in *S. sclerotiorum* only. Of particular interest was cluster b, including genes upregulated during the colonization of *A. thaliana* and on camalexin in *S. sclerotiorum.* With very few exceptions, orthologs of cluster b genes in *S. trifoliorum* were not upregulated by camalexin. We identified 70 genes of *S. sclerotiorum* upregulated on camalexin and during the colonization of *A. thaliana*, among which only 3 had orthologues induced on camalexin in *S. trifoliorum* (**Figure 6B, Supplementary Table 15**). This observation suggested that *S. sclerotiorum* but not *S. trifoliorum* is able to reprogram its transcriptome in response to camalexin. For instance, a portion of the SPREx cluster 15.5 including *S. sclerotiorum* genes *Sscle15g106410* to *Sscle15g106510* were expressed similarly during the colonization of *A. thaliana* and on camalexin (Pearson correlation coefficient ρ=0.96, adj. p-value 1.8E^-10^), while the orthologous genes in *S. trifoliorum* Scltri_004387 to Scltri_004377 were expressed very weakly on camalexin (ρ=0.19, adj. p-value 0.44) (**Figure 6C**).

**Figure 6.**
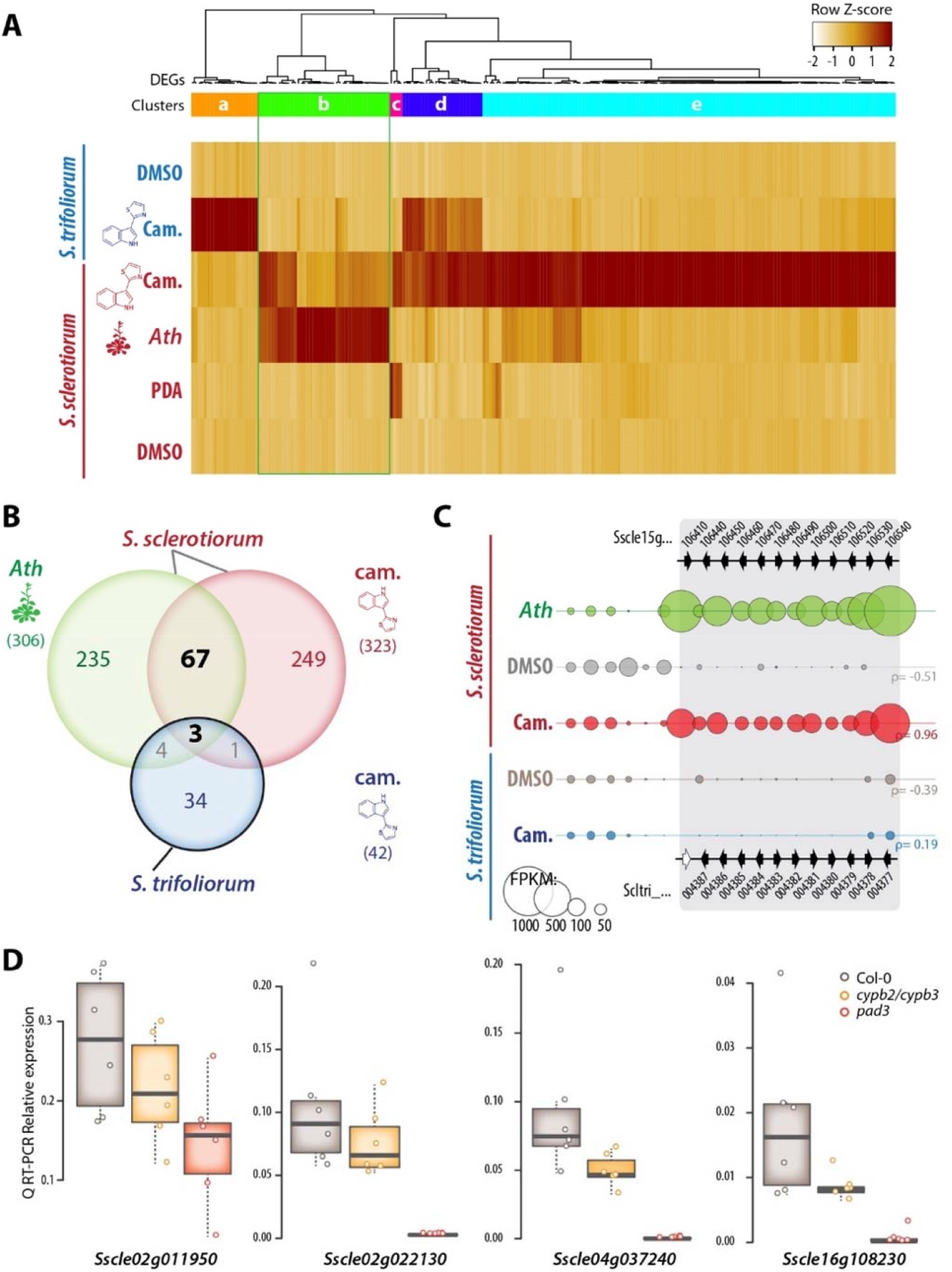
Transcriptome reprogramming in response to camalexin in *S. sclerotiorum* and *S. trifoliorum.* **(A)** Normalized FPKM expression values for differentially expressed genes during the colonization of *A. thaliana (Ath, S. sclerotiorum* only) and upon camalexin treatment (Cam., *S. sclerotiorum* and *S. trifoliorum).* Expression of orthologs of DEGs from the other *Sclerotinia* species are shown for comparison purposes. DMSO, dimethyl sulfoxide; PDA, potato dextrose agar. **(B)** Venn diagram illustrating the number of DEGs in *S. sclerotiorum* during the colonization *A. thaliana, S. sclerotiorum* growth on camalexin and orthologs of *S. trifoliorum* DEGs during growth on camalexin. Number between brackets corresponds to complete gene sets. **(C)** Expression of a subset of genes from the SPREx cluster 15.5 in *S. sclerotiorum* and the syntenic region in *S. trifoliorum* genome. The position and orientation of genes is indicated by arrowheads, labelled with gene names, empty arrowheads indicate absent genes. The size of bubbles shows the average FPKM values for each gene in five conditions. **(D)** Relative expression at 72 hours post inoculation for five *S. sclerotiorum* genes determined by quantitative reverse transcription PCR (Q RT-PCR) on *A. thaliana* wild type plants, *cyp79b2 cyp79b3* and *pad3* mutants. Values shown are for 6 independent biological replicates averaged over two technical replicates.

We identified 70 genes in *S. sclerotiorum* upregulated on camalexin and during the colonization of *A. thaliana*, indicating that these genes could respond to camalexin *in planta* (**Supplementary Table 15**). The most abundant gene ontology (GO) annotations in these genes included “no GO terms” (22), “oxidation-reduction process” (13), “carbohydrate metabolic process” (7), and “transmembrane transporter activity” (5). The most abundant PFAM annotations were “sugar (and other) transporter” (PF00083, 5), “cytochrome P450” (PF00067, 4), and “carboxylesterase family” (PF00135, 3). To confirm that camalexin and iGLs produced by *A. thaliana* are required to trigger the induction of *S. sclerotiorum* genes, we compared by quantitative RT-PCR the expression of seven *S. sclerotiorum* genes during the colonization of wild type, *cypb2/b3*, and *pad3* plants, defective in camalexin biosynthesis (**Supplementary Figure 3**). At 72 hpi, the expression of *Sscle07g055350* and *Sscle15g106410* was not different during infection of *A. thaliana* wild type and mutant plants. The expression of *Sscle02g011950* and *Sscle02g022130* was strongly reduced during infection of *cypb2/b3* but not *pad3* as compared to wild type (**Figure 6D**). The expression of *Sscle04g037240, Sscle08g067130* and *Sscle16g108230* was significantly reduced both during infection of *cypb2/b3* and *pad3* mutants as compared to wild type (Welch’s t test p-value < 0.05) (**Figure 6D**). These results indicated that host-derived cam. and iGLs modulate the expression of *S. sclerotiorum* genes during infection.

### *Cis*-regulatory variation contributed to the evolution of camalexin responsiveness in the *Sclerotinia* genus

We propose that transcriptome plasticity in host responsive genes contributed to host range variation in the *Sclerotiniaceae.* In particular, regulatory variation in the 70 *S. sclerotiorum* genes up-regulated both on *A. thaliana* and on camalexin *in vitro* (**Figure 6**, **Supplementary Table 15**) likely facilitated the colonization of plants from the *Brassicaceae.* To support this hypothesis, we first analyzed the conservation of these 70 genes in 670 species across the fungal kingdom (**Supplementary Figure 4**). Only six genes with no predicted function were detected in less than 100 fungal species, suggesting that gene presence/absence polymorphisms played a limited role in the evolution of responsiveness to *A. thaliana* and camalexin in *S. sclerotiorum.* Second, we analyzed the *cis* elements in the 1000-bp upstream sequences of these genes using the MEME-suite. The WWCCCCRC motif was significantly enriched in these 70 genes (221 sites in 65 upstream sequences, E value 0.000083). By contrast, WWCCCCRC was not significantly enriched in the upstream sequences of the 69 orthologous *S. trifoliorum* genes (41 sites in 41 upstream sequences, E value 0.58). The average distance was −480.7 bp from the start position of the ORF (**Figure 7A**). Using published yeast protein-DNA ChIP data (YEASTRACT_20130918) through TOMTOM we found five proteins of *Saccharomyces cerevisiae*, Mig1, Mig2, Mig3, Adr1, and Rsf2, that are capable of binding WWCCCCRC-like motifs. Using homology searches and phylogenetic analyses, we identified potential orthologous proteins in *S. sclerotiorum* and *S. trifoliorum* (**Supplementary Figure 5**). Probable orthologs of yeast *Mig1-3 (Multicopy Inhibitor of GAL gene expression)* and Aspergillus *CreA (Carbon catabolite repressor A)* encoding a protein harboring a central zinc finger H2C2 domain were *Sscle01g002690* and *Scltri_002966* (**Figure 7B**). This gene was slightly repressed by camalexin treatment in *S. trifoliorum* (LFC −0.25), and slightly induced by camalexin (LFC 0.60) and during *A. thaliana* colonization (LFC 0.38) in *S. sclerotiorum* (**Figure 7C**). Probable orthologs of yeast *ADR1 (Alcohol Dehydrogenase II synthesis Regulator)* and *RSF2 (Respiration factor 2)* encoding a protein harboring a N-terminal zinc finger H2C2 domain and a C-terminal fungal specific transcription factor domain PF04082 were *Sscle07g055670* and *Scltri_004270* (**Figure 7B**). This gene was slightly repressed by camalexin in *S. trifoliorum* (LFC −0.41) whereas it was induced by camalexin (LFC 0.51) and during *A. thaliana* colonization (LFC 1.58) in *S. sclerotiorum* (**Figure 7C**). These transcription factors are prime candidates for mediating large scale transcriptome reprogramming in response to camalexin in the *Sclerotinia* lineage. These results suggest that *cis*-regulatory variation in targets of SsCreA and SsADR1 contributed to the evolution of transcriptional responsiveness to camalexin in *S. sclerotiorum.*

**Figure 7.**
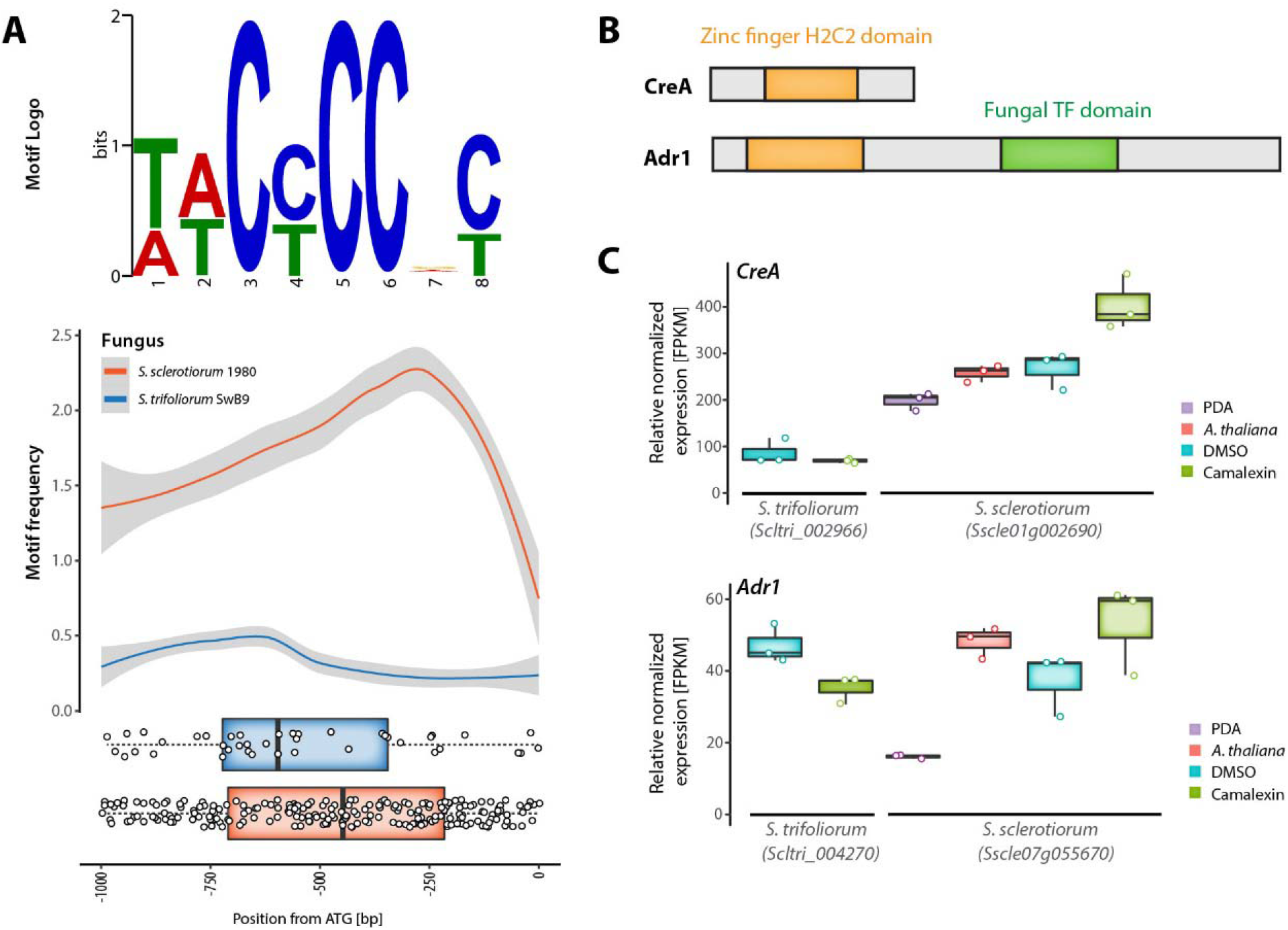
*Cis*-regulatory variation in *S. sclerotiorum* genes induced by camalexin and during the colonization of *A. thaliana.* **(A)** Top: sequence logo of the WWCCCCRC *cis*-regulatory element enriched in *S. sclerotiorum* genes induced by camalexin and *A. thaliana* infection by comparison to their *S. trifoliorum* orthologs. Bottom: distribution of the WWCCCCRC element in the 1 kbp upstream sequence of *S. sclerotiorum* genes induced by camalexin and *A. thaliana* infection and their *S. trifoliorum* orthologs. **(B)** Domain structure of *S. sclerotiorum* orthologs of CreA and Adr1 transcription factors known to bind the WWCCCCRC *cis* element in yeast. Length of the boxes is proportional to number of amino acids. **(C)** Relative normalized expression of *S. trifoliorum* and *S. sclerotiorum CreA* and *Adr1* transcription factors.

## Discussion

In this work, we analyzed the global transcriptome of *S. sclerotiorum* during the colonization of hosts from six botanical families, providing a unique opportunity to test for the existence of generalist and host-specific transcriptomes in this fungal pathogen. While previous investigations revealed adaptations to a true generalist lifestyle supporting the colonization of any host (Badet et al., 2017; Peyraud et al., 2019), we provide here molecular evidence for poly-specialism, the use of multiple independent modules dedicated to the colonization of specific hosts. These findings indicate that adaptation to new hosts can select for generalist and poly-specialist features within a single genome. We highlight a key role of regulatory variation in conserved fungal genes for the expansion of host range in this pathogen lineage.

We identified a subset of host species triggering specialized transcriptome reprogramming in *S. sclerotiorum.* Genes related to detoxification of host defense compounds were enriched in the specialized transcriptomes, while the core transcriptome overrepresented functions associated with carbohydrate catabolism and sugar transport. Host-specific regulation of pathogen genes confers the ability to quantitatively modulate the virulence program to match requirements for specific hosts. Transcriptional plasticity contributes to successful colonization of different host plants in aphids and a range of hemi-biotrophic fungal and oomycete pathogens (Mathers et al., 2017; Petre et al., 2020). Host-specialized transcriptomes are often small and consist of secreted proteins with roles in modulating host plant defense responses and nutrient assimilation such as effectors, proteases, oxidoreductases, detoxifying enzymes, and transporters (Petre et al., 2020). For example, the head blight pathogen *Fusarium graminearum* differentially expresses proteins related to nutrient transport and chitin metabolism on various host plants, i.e. wheat *(Triticum aestivum)*, barley *(Hordeum vulgare)*, and maize (*Zea mays)* (Harris et al., 2016; Lysøe et al., 2011). The septoria leaf blotch pathogen *Zymoseptoria tritici* specifically regulates oxidoreductases and detoxification-related enzymes such as cytochrome p450 proteins on wheat and brome *(Brachypodium distachyon)* (Kellner et al., 2014). A host-specialized transcriptome has been suggested previously for *S. sclerotiorum* when infecting *B. napus* and lupin *(Lupinus angustifolius)*, although the majority of induced genes were induced on both hosts (Allan et al., 2019). Our analysis on six dicot host species revealed that 52% of *S. sclerotiorum* genes upregulated *in planta* were host-specific. Since the number of host-specific induced genes varied considerably according to host (from 0% on *S. lycopersicum* to 36.9% on *B. vulgaris)*, predicting how the relative proportion of host-specific transcripts would vary if more host species were included in the analysis remains challenging. Future attempts at estimating the breadth of the *S. sclerotiorum* core transcriptome may focus on very distant plant lineages and naïve plant lineages that are susceptible to *S. sclerotiorum* but have not co-evolved with it. Such efforts would help identify a minimal virulence gene set and test whether these genes harbor evolutionary signatures distinct from host-specific transcriptome genes. Clear host-specific gene regulation suggests that transcriptome-based reverse ecology approaches are feasible, allowing for instance to identify new host defense mechanisms based on pathogen transcriptome data.

We found that the *S. sclerotiorum* core and host-specialized transcriptomes are organized in genomic clusters. Twenty-five of the 58 *Sclerotinia* plant-response co-expression (SPREx) clusters overlapped at least partly with predicted secondary metabolite biosynthetic clusters identified with AntiSMASH (Graham-Taylor et al., 2020). Metabolic gene clusters are widely found in filamentous fungi and are often associated with virulence (Wisecaver and Rokas, 2015). Like *Botrytis cinerea*, which engages in metabolic warfare with *A. thaliana* (Zhang et al., 2019c), the presence of at least 25 metabolic gene clusters that are induced on at least one host plant indicates that *S. sclerotiorum* employs various mycotoxins to engage in the metabolic battle with its host plants. Intriguingly, these mycotoxins can be deployed in a species-specific manner, exemplified with the botcinic acid gene cluster only upregulated on *A. thaliana* and upon camalexin in our dataset. In this model, *S. sclerotiorum* gains the upper hand during infection of its host plants due to effective detoxification of their metabolic compounds, for instance isothiocyanate detoxification by SsSaxA.

Gene losses and gene gains by recombination, horizontal gene transfer, or copy number variation are efficient means to modify the host range (Gluck-Thaler and Slot, 2015; Menardo et al., 2016). For instance, horizontal gene transfer contributed to host range expansion of the *Metarhizium* genus (Zhang et al., 2019a). Also, gene copy number variation is often driven by repetitive and transposable elements and contributes to pathogenicity and can provide high variability of diverging paralogs of effectors, for example in powdery mildew fungi (Frantzeskakis et al., 2018). These mechanisms can enable rapid adaptation to new hosts and contribute to host range expansion in some lineages. Nevertheless, gene presence/absence polymorphism and coding sequence changes can have detrimental effects, in particular in unstable environments. For instance, pathogen gene loss may be advantageous on host carrying a matching R protein, but detrimental otherwise. Because coding but not *cis*-regulatory mutations are sensitive to frameshift, coding sequence variation is more likely to be detrimental and pleiotropic. In addition, transcription factor binding sites are short in comparison with coding sequence and therefore more likely to be neofunctionalized. This is the reasoning behind the *cis*-regulatory hypothesis stating that mutations that alter the regulation of gene expression are more likely to contribute to phenotypic evolution (Stern and Orgogozo, 2008). In agreement with this, we provide evidence that *cis*-regulatory variation contributes to the evolution of camalexin responsiveness in *Sclerotinia. S. trifoliorum* is closely related to *S. sclerotiorum* but unable to infect plants outside of the *Fabaceae* family (Navaud et al., 2018), yet we report that the genomes of the two fungi exhibit a high level of similarity. *S. trifoliorum* is, however, highly sensitive to phytoalexins and fails to reprogram its transcriptome in response to camalexin. Our transcriptomic analysis on camalexin focused on the respective highest tolerable concentration for each of the two fungi in order to ensure a comparable physiological state amid camalexin pressure. Future analysis could focus on concentration-dependent transcriptomes of both fungi and include early stages of camalexin pressures. The inability of *S. trifoliorum* to colonize *Brassicaceae* can be in part explained by the increased susceptibility to camalexin as well as the lack of transcriptional response to the compound. Indeed *A. thaliana cyp79b2/b3* lacking iGLs (Chen et al., 2020; Stotz et al., 2011) allowed occasional colonization by the otherwise incompatible *S. trifoliorum* in our experiments. This is reminiscent of *Botrytis* species of variable host ranges, which share the vast majority of orthologous genes, while presence-absence polymorphisms were mainly observed for key secondary metabolite enzyme-encoding genes (Valero-Jiménez et al., 2019). Our findings highlight regulatory variation as an adaptive strategy for fungal pathogens jumping to new host plants. Importantly, this implies that the gene pool of ancestral pathogens is pre-equipped to adapt on non-host plants. In this scenario, host range expansion does not require the acquisition of new genes from horizontal gene transfer or outcrossing. Transcriptional plasticity induced by non-host plants may lead to the fixation of alleles with an adaptive expression pattern. In the context of the impact of plant pathogens on food security, this observation calls for monitoring not only crop pathogen populations but also their close relatives infecting wild species in the same environment.

The current study focused on *S. sclerotiorum* infection of *A. thaliana* and relationship to its typical secondary metabolites, indole glucosinolates and camalexin, owing to the tractability of this system. *S. sclerotiorum* also exhibited clear host-specific transcriptomes on castor bean, sugar beet, and common bean. On castor bean and sugar beet, genes related to the detoxification of host compounds were enriched in the specific transcriptomes. This may reflect the specific response to phytotoxins common to these host plants, such as the red beet antimicrobial phenolic secondary metabolites known as betalains (Kumar and Brooks, 2018) and the infamous ribosome-inactivating protein ricin from castor bean (Polito et al., 2019). By contrast, tomato, sunflower, and common bean induced limited host-specific transcriptomes. Flavonoids are common antimicrobial phytotoxins produced by many vascular plants and abundant in sunflower and beans for example (Weston and Mathesius, 2013). While there are more than 10,000 flavonoid structures, *S. sclerotiorum* may use a conserved pathway to tackle these toxins. Indeed, the quercetin dioxygenase SsQDO (Sscle07g059700) catalyzes the cleavage of the flavonol carbon skeleton, thus targeting a range of flavonoids (Chen et al., 2019). This gene was strongly induced on sunflower, common bean and castor bean in our dataset.

The motif WWCCCCRC was enriched in the *cis* elements of the 70 co-upregulated genes on *A. thaliana* and camalexin in *S. sclerotiorum*, while it was less abundant in the orthologous *cis* elements in *S. trifoliorum.* In baker’s yeast, the motif is recognized by the zinc-finger transcriptional regulators ADR1 and Mig1-3, which have two likely orthologues in *Sclerotinia* and *Botrytis.* Mig1-3 are orthologous to the *Aspergillus nidulans* carbon catabolite repressor CreA, which represses genes encoding carbon hydrolytic enzymes in a pH- and carbon source-dependent manner and thus allows prioritizing usage of the preferred carbon source (Orejas et al., 1999). Furthermore, plant cell wall-degrading enzymes are regulated by CreA (de Vries and Visser, 2001) and CreA is involved in mycosis disease progression due to *A. fumigatus* infection in humans (Beattie et al., 2017). *A. flavus creA* mutants are defective in aflatoxin biosynthesis and crop colonization (Fasoyin et al., 2018). In the apple blue mold pathogen *Penicillium expansum*, CreA acts as positive regulator of the mycotoxins patulin and citrinin, and *P. expansum creA* mutants are near-avirulent on apples (Tannous et al., 2018). Together with LaeA and PacC, CreA is a global regulator of virulence (Tannous et al., 2018). ADR1 on the other hand is a transcriptional regulator that regulates alcohol dehydrogenase II in yeast in glucose-lacking environments (Walther and Schüller, 2001). ADR1-regulated genes include genes in carbon metabolism related to the oxidation of non-fermentable carbon sources (Young et al., 2003). The *S. sclerotiorum* orthologous zinc finger H2C2 domain protein SsCreA *(Sscle01g002690)* could therefore be involved in the tight host-specific regulation of mycotoxin biosynthesis and plant cell wall-degrading enzymes, while SsADR1 *(Sscle07g055670)* could regulate alcohol dehydrogenases in response to certain plant-specific carbon sources. Both genes are slightly induced by camalexin, suggesting that their specific role on *A. thaliana* and *Brassicaceae* could be to respond to the presence of glucosinolates. Downstream, SsCreA and SsADR1 would then regulate the appropriate response to the host that produces the toxin, including activating the detoxification machinery for glucosinolates. As frequently observed for transcription factors, these two genes are only slightly induced and were therefore not identified by the DE analysis. An alternative possibility is that another DNA-binding protein acquired capability of binding the WWCCCCRC motif. For example, the putative C2H2 fungal transcription factor Sscle07g059580 was induced both by camalexin and on *A. thaliana* and has little homology to *Aspergillus* and *Saccharomyces* transcription factors. It is well possible that Sscle07g059580 evolved a similar function to SsADR1 but with a specific role in the response to *Brassicaceae.* The evolution of camalexin responsiveness may therefore have involved *cis*- and *trans*-regulation. Further analysis of the transcription factors SsCreA, SsADR1, and Sscle07g059580 regarding their binding efficacy of the WWCCCCRC motif and their specific function in response to *Brassicaceae* is required to answer these questions.

Although the gene pools of *S. sclerotiorum* and *S. trifoliorum* overlap nearly perfectly, a few genes were specific of each species. *S. sclerotiorum* genes absent from the *S. trifoliorum* genome notably include three genes from the botcinic acid gene cluster, which could affect the biosynthesis (although no *bona fide* secondary metabolite enzyme was missing), the regulation and the localization of the toxin. Overall, a few p450 cytochromes induced on specific hosts, such as Sscle08g067130 and Sscle11g085970 induced on *A. thaliana*, and few small secreted proteins such as Sscle06g055310 (induced on tomato) are absent in the proteome of *S. trifoliorum*, These could be key components of the infectious program on these particular hosts and their absence may partly explain the limited host range of *S. trifoliorum.* Future studies should explore the specific interaction of *S. sclerotiorum* with host-specific resistance components such as ricin and flavonoids as well as the role of host-specific genes and genes missing in the genome of *S. trifoliorum* in virulence. Since sexual recombination is rare for *Sclerotinia* and many fungi in general, the role of horizontal gene transfer and in particular the sources of genetic material should be examined in the future. It is conceivable that the plant-fungus microbiome can be a rich source of genetic material, particularly if plant pathogens or epiphytes of different kingdoms share a specific niche (plant host). Future research should therefore also focus on characterizing the genetic resources within the microbiome associated with broad-range fungal pathogens including *Sclerotinia.*

Our findings reveal that host range expansion can arise through regulatory variation in genes conserved in related non-adapted fungal pathogen species. The genetic requirements for such an evolutionary trajectory need to be determined in order to identify nonpathogenic species with a high potential for jumping to new hosts through regulatory variation. This mechanism enables the emergence of new disease with no or limited gene flow between strains and species, and could underlie the emergence of new epidemics originating from wild plants in agricultural settings.

## Materials and Methods

### Biological material

The fungal isolates *Sclerotinia sclerotiorum* isolate 1980 (Boland and Hall, 1994) and *Sclerotinia trifoliorum* SwB9 (Vleugels et al., 2013) were used in this study. The fungi were cultivated on potato dextrose agar (PDA) at 23°C or stored on PDA at 4°C. Plants were grown in Jiffy pots under controlled conditions at 23°C, with a 9 hour light period at intensity of 120 μmol m^-2^ s^-1^ for up to five weeks. For transcriptome sequencing, we used *Arabidopsis thaliana* Col-0 genotype (accession number N1093), *Solanum lycopersicum* L. cv Ailsa Craig, *Helianthus annuus* cv XRQ, *Ricinus communis* cv Hale (accession number PI 642000) and *Beta vulgaris* L. subsp. *vulgaris* (accession number PI 355961) provided by the USDA ARS Germplasm Resources Information Network, *Phaseolus vulgaris* (accession number G19833) provided by the International Center for Tropical Agriculture (CIAT). For *Sclerotinia* infection assays, *A. thaliana* accession Col-0 and the T-DNA insertion lines *cyp79b2 cyp79b3* (Zhao et al., 2002) and *pad3-1* (Zhou et al., 1999) were used.

### RNA sequencing

RNA was collected in triplicates from *S. sclerotiorum* mycelium fragments (40 μm-100 μm) cultured in liquid potato dextrose broth (PDB) for 24 hours and *S. sclerotiorum* mycelium collected from potato dextrose agar (PDA) plates center and edge (Peyraud et al., 2019). Plants were inoculated on the adaxial surface of fully developed leaves by 0.5 cm-wide plugs of PDA (potato dextrose agar, Fluka) colonized by *S. sclerotiorum* or *S. trifoliorum.* Plants were incubated in trays closed with plastic wrap to maintain 80% humidity and placed in a Percival AR-41L3 chamber under the same day/light condition as for plant growth. The center and edge of 25 mm-wide developed necrotic lesions were isolated with a scalpel blade and immediately frozen in liquid nitrogen for RNA extraction (Sucher et al., 2020). In addition, we used samples from *S. sclerotiorum* and *S. trifoliorum* cultivated on PDA with DMSO (control) or camalexin (125 μM for *S. sclerotiorum*, 25 μM for *S. trifoliorum).* All samples were generated in biological triplicates. Total RNA from infected plant samples were extracted using NucleoSpin RNA extraction kits (Macherey-Nagel) as described in (Sucher et al., 2020). For DMSO and camalexin samples, around 100 mg of liquid nitrogen frozen samples were ground with metal beads (2.5 mm diameter) in a Retschmill apparatus (24 hertz for 2×1 min) before re-suspending in 1 mL Trizol reagent (Thermo Fisher Scientific) and incubating at room temperature for 5 min. 200 μL of chloroform was added and the samples mixed thoroughly before incubating at room temperature for 3 min. After centrifugation at 12,000 *g* (4°C) for 15 min, the upper aqueous phase was recovered and nucleic acids were precipitated by adding 2 μL GlycoBlue (Ambion) and 500 μL isopropanol (incubated at −20°C for 10 min). After centrifugation at ~15,000 *g* (4°C) for 15 min, pellets were washed twice with 70% ethanol before drying and re-suspended in RNase-free water. To eliminate chloroform traces, water re-suspended nucleic acids were further cleaned using an RNA extraction kit (Qiagen, Germany) following manufacturer’s instructions. Genomic DNA was removed via DNase treatment (TURBO DNase; Ambion) following manufacturer’s instructions. Quality and concentrations of RNA were assessed with Agilent bioanalyzer nano chips. RNA sequencing (RNA-seq) was performed by Fasteris (Switzerland, Planles-Ouates) to produce Illumina reads (125 bp, paired-end) on a HiSeq2500 sequencer.

### Gene expression quantification and differential expression analyses

Trimmomatic v0.36 was employed to trim raw reads with settings for single- or paired-end sequencing and the following settings for paired-end: “ILLUMINACLIP:TruSeq2-PE.fa:5:30:10 SLIDINGWINDOW:3:18 LEADING:6 TRAILING:6 MINLEN:90” (Bolger et al., 2014). FastQC v0.11.5 (Brabham bioinformatics) was used to perform quality control of the trimmed reads. Mapping was conducted with HISAT2 (Kim et al., 2015) with “--max-intronlen 500 −k 1” against the *S. sclerotiorum* isolate 1980 reference genome (Derbyshire et al., 2017). Between 5,830,207 and 30,962,640 reads mapped uniquely to the annotated reference genome (**Supplementary Table 13**).Sorted BAM files and indices were generated with samtools v1.7 (Li et al., 2009). FPKM (fragments per kilobase of transcript per million mapped reads) tables were generated using the Cufflinks function cuffnorm with “--compatible-hits-norm --library-norm-method classic-fpkm” (Trapnell et al., 2010) (**Supplementary Table 2** and **14**). Differential expression analysis was run on 7,423 protein coding genes under a limma-edgeR pipeline (Law et al., 2016) with cut-offs of p ≤ 0.01 and log2 fold change ≤ −1 (downregulated) or ≥ 1 (upregulated) using *S. sclerotiorum* gene expression on PDA as a reference (**Supplementary Table 2** and **14**).

### Gene Ontology and PFAM domain analyses

We used Blast2GO v5.0.21 (Conesa et al., 2005; Götz et al., 2008) to generate GO and PFAM tables for both *S. sclerotiorum* 1980 (**Supplementary Table 1**) and *S. trifoliorum* SwB9 (**Supplementary Table 11**). For enrichment analyses, we considered GO and PFAM annotations present in a least one upregulated gene on each host separately. We performed chi-squared tests in R using the total number of *S. sclerotiorum* non-redundant GO (17,291) and PFAM (11,746) annotations as reference sets. P-values were adjusted using Bonferroni correction for multiple testing.

Enrichment fold for annotation ‘x’ corresponded to (Up_x_/Expressed_x_) / (Up_ALL_/Expressed_ALL_) where Up_x_ is the number of upregulated genes with annotation x, Expressed_x_ is the number of expressed genes with annotation x, Up_ALL_ is the total number of annotated genes upregulated and ExpressedALL is the total number of annotated genes. Annotations with enrichment fold > 1 and adjusted p-val < 0.01 were considered enriched with upregulated genes.

### Genomic clusters analysis

*Sclerotinia* plant-response co-expression (SPREx) clusters were identified using a sliding-window approach. The 7,523 *S. sclerotiorum* expressed genes were ordered according to their position on chromosomes. For each gene, Pearson Product-Moment correlation of expression (FPKM values) was calculated against each of the four previous and four following expressed genes on chromosomes. We calculated the average of these eight Pearson correlation coefficients to determine expression correlation within a 9-gene windows. To reduce bias due to extreme correlation values in the detection of expression clusters and to represent local expression correlation in a Circos plot, we used moving average over 20 consecutive windows. To be part of a clusters, genes had to be less than 3 genes apart from a 9-gene window with expression correlation ≥ 0.15 and expression correlation moving average ≥ 0.15. Genes located at the end of chromosomes were manually inspected and draft clusters located less than 3 genes apart were merged by manual curation. The inter-host expression variance correspond to the log_10_ of variance for FPKM values in edge samples collected on the six hosts (18 samples). To improve legibility, variance was represented as moving average over 20 consecutive genes in Circos. SPREx clusters were considered overlapping with secondary metabolite biosynthesis clusters when at least one gene was shared with clusters identified by AntiSMASH (Medema et al., 2011) reported in (Graham-Taylor et al., 2020). Whole genome data visualization was done with Circos 0.69-2 (Krzywinski et al., 2009) using script files provided as **Supplementary File 1**.

### *S. trifoliorum* genome assembly

High molecular weight DNA isolation was performed as described in (Kusch et al., 2020). Library preparation and sequencing were done at the GeT-PlaGe core facility, INRAE Toulouse, France, according to the “1D gDNA selecting for long reads (SQK-LSK108)” instructions. DNA was quantified at each step using the Qubit dsDNA HS assay kit (Life Technologies) and purity was assessed using a Nanodrop (Thermo Fisher Scientific). The Fragment Analyzer (AATI) high sensitivity DNA fragment analysis kit was used to determine size distribution and degradation. The purification steps were performed using AMPure XP beads (Beckman Coulter). For one flow cell, 5 μg of purified DNA was sheared at 8 kb using the megaruptor1 system (Diagenode), followed by a DNA damage repair step on 2 μg of sample. Then an END-repair, dA-tailing of double-stranded DNA fragments, and adapter ligation were performed on the library. The library was loaded onto one R9.4.1 flow cell and was sequenced on a GridION instrument at 0.05 pmol within 48 h. The Canu v1.6 (Koren et al., 2017) assembly yielded 48 contigs between 3,548,298 bp and 6,307 bp and a total genomic length of 40,161,953 bp. We did four cycles of polishing with Pilon (Walker et al., 2014) and then used Blobtools v1.0 (Laetsch and Blaxter, 2017) to identify contigs with *Sclerotiniaceae* identity, as described before (Kusch et al., 2020) (**Supplementary Figure 6**). To identify mitochondrial contigs, we used RNAweasel (http://megasun.bch.umontreal.ca/RNAweasel/) with the genetic code for yeast mitochondrial and performed BLASTN against the *S. sclerotiorum* mitochondrial genome (GenBank accession KT283062.1). The final genome assembly (**Table 1**) was subjected to repeat masking (RepeatMasker v4.0.7 (Smit et al., 2016), RepBase-20170127 prior to *ab initio* gene annotation with BRAKER2 (Hoff et al., 2016) where we included all RNA-seq data of *S. trifoliorum* SwB9 produced in this study, i.e. cultivated *in vitro* on PDA (1x), PDA with DMSO (3x), and with 25 μM camalexin (3x), and from infection of *P. vulgaris* (1x). All gene models were manually curated via Web Apollo (Lee et al., 2013).

**Table 1.**
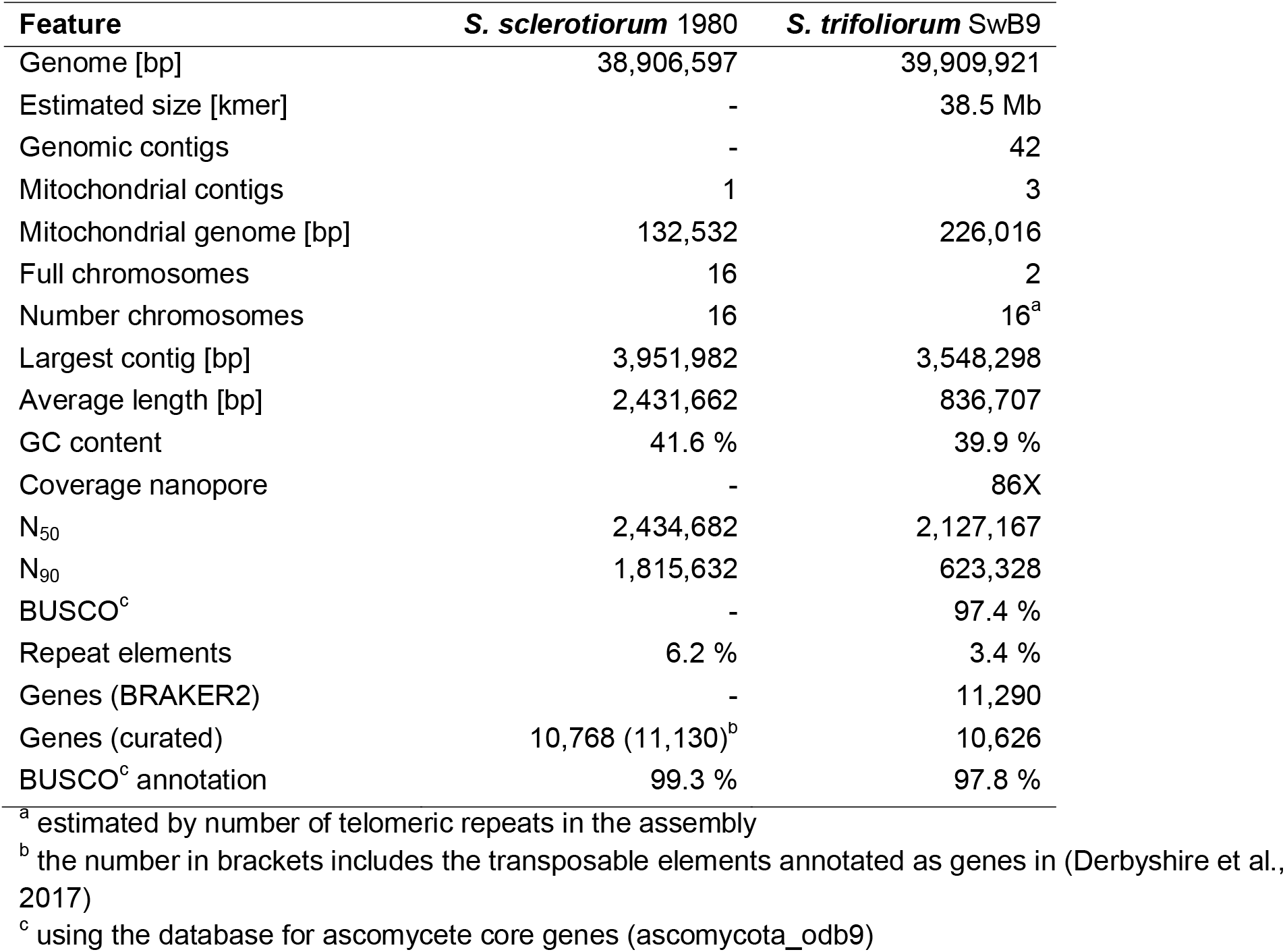
*S. trifoliorum* SwB9 genome assembly statistics.

| Feature | *S. sclerotiorum* 1980 | *S. trifoliorum* SwB9 |
| --- | --- | --- |
| Genome [bp] | 38,906,597 | 39,909,921 |
| Estimated size [kmer] | - | 38.5 Mb |
| Genomic contigs | - | 42 |
| Mitochondrial contigs | 1 | 3 |
| Mitochondrial genome [bp] | 132,532 | 226,016 |
| Full chromosomes | 16 | 2 |
| Number chromosomes | 16 | 16<sup>a</sup> |
| Largest contig [bp] | 3,951,982 | 3,548,298 |
| Average length [bp] | 2,431,662 | 836,707 |
| GC content | 41.6 % | 39.9 % |
| Coverage nanopore | - | 86X |
| N<sub>50</sub> | 2,434,682 | 2,127,167 |
| N<sub>90</sub> | 1,815,632 | 623,328 |
| BUSCO<sup>c</sup> | - | 97.4 % |
| Repeat elements | 6.2 % | 3.4 % |
| Genes (BRAKER2) | - | 11,290 |
| Genes (curated) | 10,768 (11,130)<sup>b</sup> | 10,626 |
| BUSCO<sup>c</sup> annotation | 99.3 % | 97.8 % |

<sup>a</sup> estimated by number of telomeric repeats in the assembly<sup>b</sup> the number in brackets includes the transposable elements annotated as genes in (Derbyshire et al., 2017)<sup>c</sup> using the database for ascomycete core genes (ascomycota\_odb9)

### Comparative genome and proteome analysis

The genomes of *S. sclerotiorum* Ss1980 (Derbyshire et al., 2017) and *S. trifoliorum* SwB9 were compared by synteny using MUMmer3 (Kurtz et al., 2004); synteny plotting was performed with genoPlotR (Guy et al., 2010). The proteomes of *S. sclerotiorum* Ss1980 (Derbyshire et al., 2017), *Myriosclerotinia sulcatula* MySu01 (Kusch et al., 2020), *Botrytis cinerea* B05.10 (Van Kan et al., 2017), and *S. trifoliorum* SwB9 were compared via OrthoFinder (Emms and Kelly, 2019).

### Detection of *cis* elements

The 1000-bp upstream sequences of all genes of *S. sclerotiorum* Ss1980 (Derbyshire et al., 2017) and *S. trifoliorum* SwB9 were extracted using bedtools v2.25.0 (Quinlan and Hall, 2010). The MEME-Suite 5.1.1. at http://meme-suite.org (Bailey et al., 2009, 2015) was used for motif discovery and enrichment (MEME, DREME), motif scanning (FIMO), and motif comparison (TOMTOM) against the *Saccharomyces cerevisiae* YEASTRACT_20130918.meme database. The *S. cerevisiae* motif-binding proteins were queried against the *S. sclerotiorum* Ss1980 proteome via BLASTP to identify the potential motif-binding orthologues DNA-binding proteins.

### Protein phylogenetic analysis

To identify *Sclerotinia* homologs of yeast Mig1-3, Adr1 and Rsf2, yeast protein sequences were retrieved from UniprotKB and used in BlastP search against a local database of Leotiomycete protein sequences. Best hits from *Sclerotinia* sp., *Myriosclerotinia sulcatula* and *Botrytis cinerea* species were then used in reciprocal BlastP searches against the UniprotKB database. Multiple sequence alignments were generated with Muscle and manually curated in Jalview to remove gaps and poor alignment quality regions. This resulted in alignments of 28 sequences and 217 informative positions (Mig1-3, **Supplementary File 2**) and 26 sequences and 129 informative positions (Adr1/Rsf2, **Supplementary File 2**). Maximum likelihood phylogenies were generated in phylogeny.fr using PhyML and the WAG model for amino acid substitutions using a discrete gamma model with 4 categories (**Supplementary File 2, Supplementary Figure 5**).

### Data analysis and plotting

R v.3.5.1 (R Core Team, 2018) was used for statistical analysis and generation of plots. The ggplot function from the ggplot2 library (Wickham, 2009) was used to generate the plots. Codes for data analysis are deposited at https://github.com/stefankusch/sclerotinia_2020.

## Author contributions

SK and SR designed the project. RNA-seq sampling was performed by JL, ON, MM and SK. SK and SR performed data analysis, statistical analysis, and plotting. Genome nanopore sequencing was done by CZ, CR, and CD, assembly of S*. trifoliorum* SwB9 v1 genome by SK, and gene annotation and manual curation by SK and HI. Further experiments and pathogen assays were performed by SK, SR, JL, and NG. SK and SR drafted the manuscript, all authors edited and proofread the manuscript.

## Acknowledgements

We are grateful to the genotoul bioinformatics platform Toulouse Midi-Pyrenees (Bioinfo Genotoul, http://bioinfo.genotoul.fr/) for providing help and/or computing and/or storage resources. This work was performed in collaboration with the GeT core facility, Toulouse, France (http://get.genotoul.fr), and was supported by France Génomique National infrastructure, funded as part of “Investissement d’avenir” program managed by Agence Nationale pour la Recherche (contract ANR-10-INBS-09). We thank Marielle Barascud for excellent technical assistance.

## Data availability statement

Raw RNA-seq reads data and processed gene expression files are available from the NCBI GEO under accessions GSE106811, GSE138039, and GSE116194. The new RNA-seq data generated for this study (raw reads and re-analysis of gene expression) was deposited in NCBI GEO under accession GSE159792. *S. trifoliorum* genome assembly and annotation are available from the European Nucleotide Archive (ENA) under project accession PRJEB36746 and genome accession ERZ1667436. Command line and R scripts are available at https://github.com/stefankusch/sclerotinia_2020. All other datasets analyzed for this study are included in the manuscript and the supplementary files.

## Abbreviations

DE: Differential expression
DEG: Differentially expressed gene
FPKM: Fragments per kilobase of transcript per million mapped reads
GO: Gene ontology
OG: Orthogroup
PDA: Potato dextrose agar
PDB: Potato dextrose broth
RNA-seq: Whole transcriptome shotgun sequencing
TE: Transposable element

## SUPPLEMENTARY MATERIAL

**Supplementary Figure 1.** Principal component analysis of *S. sclerotiorum* gene expression during the colonization of six plants, samples harvested at the edge of invasive colonies. PDA, Potato Dextrose Agar, *Ath, Arabidopsis thaliana; Bvu, Beta vulgaris; Han, Helianthus annuus; Pvu, Phaseolus vulgaris; Rco, Ricinus communis; Sly, Solanum lycopersicum.*

**Supplementary Figure 2.** Complements to the phytoalexin *in vitro* sensitivity assay. The assay was performed as described in Figure 5, additional concentrations and phytoalexin solutions are presented here. ICA/I3A/RA is a mixture of Indole-3 carboxylic acid (in ethanol), Indole-3 ylmethylamine (in DMSO) and raphanusamic acid (in water). Non-viable concentrations are labelled in red.

**Supplementary Figure 3.** Relative expression at 72 hours post inoculation for 3 *S. sclerotiorum* genes determined by quantitative reverse transcription PCR (Q RT-PCR) on *A. thaliana* wild type plants, *cyp79b2/cyp79b3* and *pad3* mutants. Values shown are for 6 independent biological replicates averaged over two technical replicates.

**Supplementary Figure 4.** Phylogenetic distribution of the 70 *S. sclerotiorum* genes upregulated during *A. thaliana* infection and on camalexin. Conservation of *S. sclerotiorum* genes (columns) in 629 fungal species (rows) is shows as a brown box when a BlatsP hit with p-value < 1E-10 was detected, as light grey boxes otherwise. Fungal species are organized following the species phylogeny retrieved from JGI Mycocosm, number in brackets indicate the number of species per lineage. The % of conserved genes bars show the proportion of genes detected in each species. The conservation % bars show the proportion of fungal species in which each gene was detected.

**Supplementary Figure 5.** Supplementary Figure 5. Phylogenetic analysis of *Sclerotinia* orthologs of yeast *Mig1-3* and *Adr1/Rsf2* genes. (A, C) Maximum likelihood phylogenetic trees obtained with PhyML using the WAG model for amino acid substitutions. The trees were rooted on *Schizosaccharomyces pombe* closest homolog. Labels show branch support determined by an approximate likelihood ratio test. (B, D) Multiple sequence alignments of the zinc finger regions for *S. sclerotiorum* and *Saccharomyces cereviseae* homologs. Conserved residues are colored according to % conservation and residue type.

**Supplementary Figure 6.** Blobtools analysis for *S. trifoliorum* genome assembly. The graph is showing nanopore read re-mapping coverage per contig on the Y-axis and GC content per contig on the X-axis. The size of the bubble indicates the size of the respective contig. Identity was determined via megablast against the NCBI nt database.

**Supplementary File 1**. Script, data and configuration files used for the generation of the Circos plot presented in Figure 2.

**Supplementary File 2.** Multiple sequence alignments and newick tree files from the phylogenetic analysis of *Sclerotinia* orthologs of yeast Mig1-3 and Adr1/Rsf2 genes

**Supplementary Table 1.** Blast2GO and RepeatMasker updated annotations of the *Sclerotinia sclerotiorum* 1980 proteome

**Supplementary Table 2.** FPKM values in the 45 RNA-seq libraries analyzed in this work for 7,423 *S. sclerotiorum* genes with FPKM > 25 in at least one library

**Supplementary Table 3.** Differential expression analysis on six host plants performed with limma-edgeR for 7423 *S. sclerotiorum* protein coding genes

**Supplementary Table 4.** Number of *S. sclerotiorum* genes differentially expressed on each host

**Supplementary Table 5.** Distribution of DEGs according to hosts in which they are differentially expressed

**Supplementary Table 6.** Chi-squared analysis of Gene Ontologies enriched with upregulated genes

**Supplementary Table 7.** Chi-squared analysis of PFAM domains enriched with upregulated genes

**Supplementary Table 8.** Identification of host-response co-expression clusters using a sliding window approach

**Supplementary Table 9.** Summary table of host-response co-expression clusters features

**Supplementary Table 10.** Chi-squared analysis of GOs and PFAMs enriched with genes in host-response co-expression clusters

**Supplementary Table 11.** Blast2GO and RepeatMasker annotations of the *Sclerotinia trifoliorum* SwB9 proteome

**Supplementary Table 12.** Identifiers of *S. sclerotiorum, S. trifoliorum, B. cinerea* and *M. sulcatula* genes in orthogroups

**Supplementary Table 13.** *S. sclerotiorum* and *S. trifoliorum* RNA-seq mapping statistics

**Supplementary Table 14.** FPKM values and Differential expression analysis on camalexin for 7423 *S. sclerotiorum* protein coding genes and their *S. trifoliorum* orthologs

**Supplementary Table 15.** Expression and taxonomic conservation data for the 70 *S. sclerotiorum* genes upregulated during *A. thaliana* colonization and growth on camalexin.

## References

Allan, J., Regmi, R., Denton-Giles, M., Kamphuis, L. G., and Derbyshire, M. C. (2019). The host generalist phytopathogenic fungus *Sclerotinia sclerotiorum* differentially expresses multiple metabolic enzymes on two different plant hosts. Sci. Rep. 9, 19966. doi:10.1038/s41598-019-56396-w.

Ámon, J., Fernández-Martín, R., Bokor, E., Cultrone, A., Kelly, J. M., Flipphi, M., et al. (2017). A eukaryotic nicotinate-inducible gene cluster: convergent evolution in fungi and bacteria. Open Biol. 7, 170199. doi:10.1098/rsob.170199.

Badet, T., Peyraud, R., Mbengue, M., Navaud, O., Derbyshire, M. C., Oliver, R. P., et al. (2017). Codon optimization underpins generalist parasitism in fungi. Elife 6, e22472. doi:10.7554/eLife.22472.

Bailey, T. L., Boden, M., Buske, F. A., Frith, M., Grant, C. E., Clementi, L., et al. (2009). MEME SUITE: tools for motif discovery and searching. Nucleic Acids Res. 37, W202–208. doi:10.1093/nar/gkp335.

Bailey, T. L., Johnson, J., Grant, C. E., and Noble, W. S. (2015). The MEME Suite. Nucleic Acids Res. 43, W39–49. doi:10.1093/nar/gkv416.

Baroncelli, R., Amby, D. B., Zapparata, A., Sarrocco, S., Vannacci, G., Le Floch, G., et al. (2016). Gene family expansions and contractions are associated with host range in plant pathogens of the genus *Colletotrichum*. BMC Genomics 17, 555. doi:10.1186/s12864-016-2917-6.

Beattie, S. R., Mark, K. M. K., Thammahong, A., Ries, L. N. A., Dhingra, S., Caffrey-Carr, A. K., et al. (2017). Filamentous fungal carbon catabolite repression supports metabolic plasticity and stress responses essential for disease progression. PLOS Pathog. 13, e1006340. doi:10.1371/journal.ppat.1006340.

Boland, G. J., and Hall, R. (1994). Index of plant hosts of *Sclerotinia sclerotiorum*. Can. J. Plant Pathol. 16, 93–108. doi:10.1080/07060669409500766.

Bolger, A. M., Lohse, M., and Usadel, B. (2014). Trimmomatic: a flexible trimmer for Illumina sequence data. Bioinformatics 30, 2114–2120. doi:10.1093/bioinformatics/btu170.

Cauldwell, A. V., Long, J. S., Moncorgé, O., and Barclay, W. S. (2014). Viral determinants of influenza A virus host range. J. Gen. Virol. 95, 1193–1210. doi:10.1099/vir.0.062836-0.

Chen, J., Ullah, C., Reichelt, M., Beran, F., Yang, Z.-L., Gershenzon, J., et al. (2020). The phytopathogenic fungus *Sclerotinia sclerotiorum* detoxifies plant glucosinolate hydrolysis products via an isothiocyanate hydrolase. Nat. Commun. 11, 3090. doi:10.1038/s41467-020-16921-2.

Chen, J., Ullah, C., Reichelt, M., Gershenzon, J., and Hammerbacher, A. (2019). *Sclerotinia sclerotiorum* circumvents flavonoid defenses by catabolizing flavonol glycosides and aglycones. Plant Physiol. 180, 1975–1987. doi:10.1104/pp.19.00461.

Conesa, A., Götz, S., Garcia-Gomez, J. M., Terol, J., Talon, M., and Robles, M. (2005). Blast2GO: a universal tool for annotation, visualization and analysis in functional genomics research. Bioinformatics 21, 3674–3676. doi:10.1093/bioinformatics/bti610.

Dalmais, B., Schumacher, J., Moraga, J., Le Pêcheur, P., Tudzynski, B., Collado, I. G., et al. (2011). The *Botrytis cinerea* phytotoxin botcinic acid requires two polyketide synthases for production and has a redundant role in virulence with botrydial. Mol. Plant Pathol. 12, 564–579. doi:10.1111/j.1364-3703.2010.00692.x.

de Vries, R. P., and Visser, J. (2001). *Aspergillus* enzymes involved in degradation of plant cell wall polysaccharides. Microbiol. Mol. Biol. Rev. 65, 497–522. doi:10.1128/MMBR.65.4.497-522.2001.

Derbyshire, M. C., Denton-Giles, M., Hane, J. K., Chang, S., Mousavi-Derazmahalleh, M., Raffaele, S., et al. (2019a). A whole genome scan of SNP data suggests a lack of abundant hard selective sweeps in the genome of the broad host range plant pathogenic fungus *Sclerotinia sclerotiorum*. PLoS One 14, e0214201. doi:10.1371/journal.pone.0214201.

Derbyshire, M. C., Denton-Giles, M., Hegedus, D. D., Seifbarghi, S., Rollins, J. A., Van Kan, J. A. L., et al. (2017). The complete genome sequence of the phytopathogenic fungus *Sclerotinia sclerotiorum* reveals insights into the genome architecture of broad host range pathogens. Genome Biol. Evol. 9, 593–618. doi:10.1093/gbe/evx030.

Derbyshire, M., Mbengue, M., Barascud, M., Navaud, O., and Raffaele, S. (2019b). Small RNAs from the plant pathogenic fungus *Sclerotinia sclerotiorum* highlight host candidate genes associated with quantitative disease resistance. Mol. Plant Pathol. 20, 1279–1297. doi:10.1111/mpp.12841.

Dong, S., Raffaele, S., and Kamoun, S. (2015). The two-speed genomes of filamentous pathogens: Waltz with plants. Curr. Opin. Genet. Dev. 35, 57–65. doi:10.1016/j.gde.2015.09.001.

Dong, S., Stam, R., Cano, L. M., Song, J., Sklenar, J., Yoshida, K., et al. (2014). Effector specialization in a lineage of the Irish potato famine pathogen. Science 343, 552–555. doi:10.1126/science.1246300.

Ebert, D., and Fields, P. D. (2020). Host–parasite co-evolution and its genomic signature. Nat. Rev. Genet. doi:10.1038/s41576-020-0269-1.

Emms, D. M., and Kelly, S. (2015). OrthoFinder: solving fundamental biases in whole genome comparisons dramatically improves orthogroup inference accuracy. Genome Biol. 16, 157. doi:10.1186/s13059-015-0721-2.

Emms, D. M., and Kelly, S. (2019). OrthoFinder: phylogenetic orthology inference for comparative genomics. Genome Biol. 20, 238. doi:10.1186/s13059-019-1832-y.

Fasoyin, O. E., Wang, B., Qiu, M., Han, X., Chung, K.-R., and Wang, S. (2018). Carbon catabolite repression gene *creA* regulates morphology, aflatoxin biosynthesis and virulence in *Aspergillus flavus*. Fungal Genet. Biol. 115, 41–51. doi:10.1016/j.fgb.2018.04.008.

Flor, H. H. (1971). Current status of the gene-for-gene concept. Annu. Rev. Phytopathol. 9, 275–296. doi:10.1146/annurev.py.09.090171.001423.

Frantzeskakis, L., Kracher, B., Kusch, S., Yoshikawa-Maekawa, M., Bauer, S., Pedersen, C., et al. (2018). Signatures of host specialization and a recent transposable element burst in the dynamic one-speed genome of the fungal barley powdery mildew pathogen. BMC Genomics 19, 381. doi:10.1186/s12864-018-4750-6.

Gluck-Thaler, E., and Slot, J. C. (2015). Dimensions of horizontal gene transfer in eukaryotic microbial pathogens. PLOS Pathog. 11, e1005156. doi:10.1371/journal.ppat.1005156.

Götz, S., Garcia-Gomez, J. M., Terol, J., Williams, T. D., Nagaraj, S. H., Nueda, M. J., et al. (2008). High-throughput functional annotation and data mining with the Blast2GO suite. Nucleic Acids Res. 36, 3420–3435. doi:10.1093/nar/gkn176.

Graham-Taylor, C., Kamphuis, L. G., and Derbyshire, M. C. (2020). A detailed *in silico* analysis of secondary metabolite biosynthesis clusters in the genome of the broad host range plant pathogenic fungus *Sclerotinia sclerotiorum*. BMC Genomics 21, 7. doi:10.1186/s12864-019-6424-4.

Guy, L., Roat Kultima, J., and Andersson, S. G. E. (2010). genoPlotR: comparative gene and genome visualization in R. Bioinformatics 26, 2334–2335. doi:10.1093/bioinformatics/btq413.

Guyon, K., Balagué, C., Roby, D., and Raffaele, S. (2014). Secretome analysis reveals effector candidates associated with broad host range necrotrophy in the fungal plant pathogen *Sclerotinia sclerotiorum*. BMC Genomics 15, 336. doi:10.1186/1471-2164-15-336.

Harris, L. J., Balcerzak, M., Johnston, A., Schneiderman, D., and Ouellet, T. (2016). Host-preferential *Fusarium graminearum* gene expression during infection of wheat, barley, and maize. Fungal Biol. 120, 111–123. doi:10.1016/j.funbio.2015.10.010.

Hoff, K. J., Lange, S., Lomsadze, A., Borodovsky, M., and Stanke, M. (2016). BRAKER1: Unsupervised RNA-Seq-based genome annotation with GeneMark-ET and AUGUSTUS. Bioinformatics 32, 767–769. doi:10.1093/bioinformatics/btv661.

Hu, X., Xiao, G., Zheng, P., Shang, Y., Su, Y., Zhang, X., et al. (2014). Trajectory and genomic determinants of fungal-pathogen speciation and host adaptation. Proc. Natl. Acad. Sci. U. S. A. 111, 16796–16801. doi:10.1073/pnas.1412662111.

Ibrahim, H. M. M., Kusch, S., Didelon, M., and Raffaele, S. (2020). Genome-wide alternative splicing profiling in the fungal plant pathogen *Sclerotinia sclerotiorum* during the colonization of diverse host families. Mol. Plant Pathol., 13006. doi:10.1111/mpp.13006.

Janzen, D. H. (1980). When is it coevolution? Evolution (N. Y). 34, 611–612. doi:10.1111/j.1558-5646.1980.tb04849.x.

Jones, J. D. G., and Dangl, J. L. (2006). The plant immune system. Nature 444, 323–329. doi:10.1038/nature05286.

Kellner, R., Bhattacharyya, A., Poppe, S., Hsu, T. Y., Brem, R. B., and Stukenbrock, E. H. (2014). Expression profiling of the wheat pathogen *Zymoseptoria tritici* reveals genomic patterns of transcription and host-specific regulatory programs. Genome Biol. Evol. 6, 1353–1365. doi:10.1093/gbe/evu101.

Kim, D., Langmead, B., and Salzberg, S. L. (2015). HISAT: a fast spliced aligner with low memory requirements. Nat. Methods 12, 357–360. doi:10.1038/nmeth.3317.

Koren, S., Walenz, B. P., Berlin, K., Miller, J. R., Bergman, N. H., and Phillippy, A. M. (2017). Canu: scalable and accurate long-read assembly via adaptive k-mer weighting and repeat separation. Genome Res. 27, 722–736. doi:10.1101/gr.215087.116.

Krzywinski, M., Schein, J., Birol, I., Connors, J., Gascoyne, R., Horsman, D., et al. (2009). Circos: An information aesthetic for comparative genomics. Genome Res. 19, 1639–1645. doi:10.1101/gr.092759.109.

Kumar, S., and Brooks, M. S.-L. (2018). Use of red beet *(Beta vulgaris* L.) for antimicrobial applications - a critical review. Food Bioprocess Technol. 11, 17–42. doi:10.1007/s11947-017-1942-z.

Kurtz, S., Phillippy, A., Delcher, A. L., Smoot, M., Shumway, M., Antonescu, C. M., et al. (2004). Versatile and open software for comparing large genomes. Genome Biol. 5, R12. doi:10.1186/gb-2004-5-2-r12.

Kusch, S., Ibrahim, H. M. M., Zanchetta, C., Lopez-Roques, C., Donnadieu, C., and Raffaele, S. (2020). A chromosome-scale genome assembly resource for *Myriosclerotinia sulcatula* infecting sedge grass *(Carex* sp.). Mol. Plant-Microbe Interact. MPMI 33, 880–883. doi:10.1094/MPMI-03-20-0060-A.

Laetsch, D. R., and Blaxter, M. L. (2017). BlobTools: Interrogation of genome assemblies. F1000Research 6, 1287. doi:10.12688/f1000research.12232.1.

Law, C. W., Alhamdoosh, M., Su, S., Smyth, G. K., and Ritchie, M. E. (2016). RNA-seq analysis is easy as 1-2-3 with limma, Glimma and edgeR. F1000Research 5, 1408. doi:10.12688/f1000research.9005.1.

Lee, E., Helt, G. A., Reese, J. T., Munoz-Torres, M. C., Childers, C. P., Buels, R. M., et al. (2013). Web Apollo: a web-based genomic annotation editing platform. Genome Biol. 14, R93. doi:10.1186/gb-2013-14-8-r93.

Li, H., Handsaker, B., Wysoker, A., Fennell, T., Ruan, J., Homer, N., et al. (2009). The Sequence Alignment/Map format and SAMtools. Bioinformatics 25, 2078–2079. doi:10.1093/bioinformatics/btp352.

Lysøe, E., Seong, K.-Y., and Kistler, H. C. (2011). The transcriptome of *Fusarium graminearum* during the infection of wheat. Mol. Plant-Microbe Interact. MPMI 24, 995–1000. doi:10.1094/MPMI-02-11-0038.

Mathers, T. C., Chen, Y., Kaithakottil, G., Legeai, F., Mugford, S. T., Baa-Puyoulet, P., et al. (2017). Rapid transcriptional plasticity of duplicated gene clusters enables a clonally reproducing aphid to colonise diverse plant species. Genome Biol. 18, 27. doi:10.1186/s13059-016-1145-3.

Medema, M. H., Blin, K., Cimermancic, P., de Jager, V., Zakrzewski, P., Fischbach, M. A., et al. (2011). antiSMASH: Rapid identification, annotation and analysis of secondary metabolite biosynthesis gene clusters in bacterial and fungal genome sequences. Nucleic Acids Res. 39, W339–W346. doi:10.1093/nar/gkr466.

Menardo, F., Praz, C. R., Wyder, S., Ben-David, R., Bourras, S., Matsumae, H., et al. (2016). Hybridization of powdery mildew strains gives rise to pathogens on novel agricultural crop species. Nat. Genet. 48, 201–205. doi:10.1038/ng.3485.

Möller, M., and Stukenbrock, E. H. (2017). Evolution and genome architecture in fungal plant pathogens. Nat. Rev. Microbiol. doi:10.1038/nrmicro.2017.76.

Navaud, O., Barbacci, A., Taylor, A., Clarkson, J. P., and Raffaele, S. (2018). Shifts in diversification rates and host jump frequencies shaped the diversity of host range among *Sclerotiniaceae* fungal plant pathogens. Mol. Ecol. 27, 1309–1323. doi:10.1111/mec.14523.

Nylin, S., and Janz, N. (2009). Butterfly host plant range: an example of plasticity as a promoter of speciation? Evol. Ecol. 23, 137–146. doi:10.1007/s10682-007-9205-5.

Orejas, M., MacCabe, A. P., Perez Gonzalez, J. A., Kumar, S., and Ramon, D. (1999). Carbon catabolite repression of the *Aspergillus nidulans xlnA* gene. Mol. Microbiol. 31, 177–184. doi:10.1046/j.1365-2958.1999.01157.x.

Otto, T. D., Gilabert, A., Crellen, T., Böhme, U., Arnathau, C., Sanders, M., et al. (2018). Genomes of all known members of a *Plasmodium* subgenus reveal paths to virulent human malaria. Nat. Microbiol. 3, 687–697. doi:10.1038/s41564-018-0162-2.

Pedras, M. S. C., Ahiahonu, P. W. K., and Hossain, M. (2004). Detoxification of the cruciferous phytoalexin brassinin in *Sclerotinia sclerotiorum* requires an inducible glucosyltransferase. Phytochemistry 65, 2685–2694. doi:10.1016/j.phytochem.2004.08.033.

Petre, B., Lorrain, C., Stukenbrock, E. H., and Duplessis, S. (2020). Host-specialized transcriptome of plant-associated organisms. Curr. Opin. Plant Biol. 56, 81–88. doi:10.1016/j.pbi.2020.04.007.

Peyraud, R., Mbengue, M., Barbacci, A., and Raffaele, S. (2019). Intercellular cooperation in a fungal plant pathogen facilitates host colonization. Proc. Natl. Acad. Sci. U. S. A. 116, 3193–3201. doi:10.1073/pnas.1811267116.

Polito, L., Bortolotti, M., Battelli, M. G., Calafato, G., and Bolognesi, A. (2019). Ricin: An ancient story for a timeless plant toxin. Toxins (Basel). 11, 324. doi:10.3390/toxins11060324.

Quinlan, A. R., and Hall, I. M. (2010). BEDTools: a flexible suite of utilities for comparing genomic features. Bioinformatics 26, 841–842. doi:10.1093/bioinformatics/btq033.

R Core Team (2018). R: A language and environment for statistical computing. R Found. Stat. Comput. Vienna, Austria. Available at: http://www.r-project.org/.

Raffaele, S., Farrer, R. A., Cano, L. M., Studholme, D. J., MacLean, D., Thines, M., et al. (2010). Genome evolution following host jumps in the Irish potato famine pathogen lineage. Science (80-.). 330, 1540–1543. doi:10.1126/science.1193070.

Raffaele, S., and Kamoun, S. (2012). Genome evolution in filamentous plant pathogens: Why bigger can be better. Nat. Rev. Microbiol. 10, 417–430. doi:10.1038/nrmicro2790.

Sánchez-Vallet, A., Fouché, S., Fudal, I., Hartmann, F. E., Soyer, J. L., Tellier, A., et al. (2018). The genome biology of effector gene evolution in filamentous plant pathogens. Annu. Rev. Phytopathol. 56, 21–40. doi:10.1146/annurev-phyto-080516-035303.

Sexton, A. C., Minic, Z., Cozijnsen, A. J., Pedras, M. S. C., and Howlett, B. J. (2009). Cloning, purification and characterisation of brassinin glucosyltransferase, a phytoalexin-detoxifying enzyme from the plant pathogen *Sclerotinia sclerotiorum*. Fungal Genet. Biol. 46, 201–209. doi:10.1016/j.fgb.2008.10.014.

Simão, F. A., Waterhouse, R. M., Ioannidis, P., Kriventseva, E. V., and Zdobnov, E. M. (2015). BUSCO: assessing genome assembly and annotation completeness with single-copy orthologs. Bioinformatics 31, 3210–3212. doi:10.1093/bioinformatics/btv351.

Simon, J.-C., D’Alencon, E., Guy, E., Jacquin-Joly, E., Jaquiery, J., Nouhaud, P., et al. (2015). Genomics of adaptation to host-plants in herbivorous insects. Brief. Funct. Genomics 14, 413–423. doi:10.1093/bfgp/elv015.

Smit, A. F. A., Hubley, R., and Green, P. (2016). Masker Open-4.0. 2013-2015. Available at: http://www.repeatmasker.org.

Sperschneider, J., Dodds, P. N., Gardiner, D. M., Singh, K. B., and Taylor, J. M. (2018). Improved prediction of fungal effector proteins from secretomes with EffectorP 2.0. Mol. Plant Pathol. 19, 2094–2110. doi:10.1111/mpp.12682.

Stern, D. L., and Orgogozo, V. (2008). The loci of evolution: How predictable is genetic evolution? Evolution (N. Y). 62, 2155–2177. doi:10.1111/j.1558-5646.2008.00450.x.

Stotz, H. U., Sawada, Y., Shimada, Y., Hirai, M. Y., Sasaki, E., Krischke, M., et al. (2011). Role of camalexin, indole glucosinolates, and side chain modification of glucosinolate-derived isothiocyanates in defense of *Arabidopsis* against *Sclerotinia sclerotiorum*. Plant J. 67, 81–93. doi:10.1111/j.1365-313X.2011.04578.x.

Sucher, J., Mbengue, M., Dresen, A., Barascud, M., Didelon, M., Barbacci, A., et al. (2020). Phylotranscriptomics of the Pentapetalae reveals frequent regulatory variation in plant local responses to the fungal pathogen *Sclerotinia sclerotiorum*. Plant Cell 32, 1820–1844. doi:10.1105/tpc.19.00806.

Tannous, J., Kumar, D., Sela, N., Sionov, E., Prusky, D., and Keller, N. P. (2018). Fungal attack and host defence pathways unveiled in near-avirulent interactions of *Penicillium expansum creA* mutants on apples. Mol. Plant Pathol. 19, 2635–2650. doi:10.1111/mpp.12734.

Thines, M. (2019). An evolutionary framework for host shifts – jumping ships for survival. New Phytol. 224, 605–617. doi:10.1111/nph.16092.

Thrall, P. H., Laine, A.-L., Ravensdale, M., Nemri, A., Dodds, P. N., Barrett, L. G., et al. (2012). Rapid genetic change underpins antagonistic coevolution in a natural host-pathogen metapopulation. Ecol. Lett. 15, 425–435. doi:10.1111/j.1461-0248.2012.01749.x.

Trapnell, C., Williams, B. A., Pertea, G. M., Mortazavi, A., Kwan, G., van Baren, M. J., et al. (2010). Transcript assembly and quantification by RNA-Seq reveals unannotated transcripts and isoform switching during cell differentiation. Nat. Biotechnol. 28, 511–515. doi:10.1038/nbt.1621.

Troch, V., Audenaert, K., Wyand, R. A., Haesaert, G., Höfte, M., and Brown, J. K. M. (2014). *Formae speciales* of cereal powdery mildew: close or distant relatives? Mol. Plant Pathol. 15, 304–314. doi:10.1111/mpp.12093.

Valero-Jiménez, C. A., Veloso, J., Staats, M., and van Kan, J. A. L. (2019). Comparative genomics of plant pathogenic *Botrytis* species with distinct host specificity. BMC Genomics 20, 203. doi:10.1186/s12864-019-5580-x.

Van Kan, J. A. L., Stassen, J. H. M., Mosbach, A., Van Der Lee, T. A. J., Faino, L., Farmer, A. D., et al. (2017). A gapless genome sequence of the fungus *Botrytis cinerea*. Mol. Plant Pathol. 18, 75–89. doi:10.1111/mpp.12384.

Vleugels, T., Baert, J., and van Bockstaele, E. (2013). Morphological and pathogenic characterization of genetically diverse *Sclerotinia* isolates from European red clover crops *(Trifolium pratense* L.). J. Phytopathol. 161, 254–262. doi:10.1111/jph.12056.

Walker, B. J., Abeel, T., Shea, T., Priest, M., Abouelliel, A., Sakthikumar, S., et al. (2014). Pilon: An integrated tool for comprehensive microbial variant detection and genome assembly improvement. PLoS One 9, e112963. doi:10.1371/journal.pone.0112963.

Walther, K., and Schüller, H.-J. (2001). Adr1 and Cat8 synergistically activate the glucose-regulated alcohol dehydrogenase gene ADH2 of the yeast *Saccharomyces cerevisiae*. Microbiology 147, 2037–2044. doi:10.1099/00221287-147-8-2037.

West-Eberhard, M. J. (1989). Phenotypic plasticity and the origins of diversity. Annu. Rev. Ecol. Syst. 20, 249–278. doi:10.1146/annurev.es.20.110189.001341.

Weston, L. A., and Mathesius, U. (2013). Flavonoids: Their structure, biosynthesis and role in the rhizosphere, including allelopathy. J. Chem. Ecol. 39, 283–297. doi:10.1007/s10886-013-0248-5.

Wickham, H. (2009). ggplot2: Elegant Graphics for Data Analysis. 1st ed., eds. R. Gentleman, K. Hornik, and G. Parmigiani New York: Springer-Verlag. doi:10.1007/978-0-387-98141-3.

Wisecaver, J. H., and Rokas, A. (2015). Fungal metabolic gene clusters - caravans traveling across genomes and environments. Front. Microbiol. 6, 161. doi:10.3389/fmicb.2015.00161.

Wyka, S. A., Mondo, S. J., Miao Liu, J. D., Nalam, V., and Broders, K. D. (2020). Whole genome comparisons of ergot fungi reveals the divergence and evolution of species within the genus *Claviceps* are the result of varying mechanisms driving genome evolution and host range expansion. bioRxiv, 2020.04.13.039230. doi:doi.org/10.1101/2020.04.13.039230.

Young, E. T., Dombek, K. M., Tachibana, C., and Ideker, T. (2003). Multiple pathways are co-regulated by the protein kinase Snf1 and the transcription factors Adr1 and Cat8. J. Biol. Chem. 278, 26146–26158. doi:10.1074/jbc.M301981200.

Zhang, Q., Chen, X., Xu, C., Zhao, H., Zhang, X., Zeng, G., et al. (2019a). Horizontal gene transfer allowed the emergence of broad host range entomopathogens. Proc. Natl. Acad. Sci. U. S. A. 116, 7982–7989. doi:10.1073/pnas.1816430116.

Zhang, S., Gu, S., Ni, X., and Li, X. (2019b). Genome size reversely correlates with host plant range in *Helicoverpa* species. Front. Physiol. 10, 29. doi:10.3389/fphys.2019.00029.

Zhang, W., Corwin, J. A., Copeland, D. H., Feusier, J., Eshbaugh, R., Cook, D. E., et al. (2019c). Plant-necrotroph co-transcriptome networks illuminate a metabolic battlefield. Elife 8, e44279. doi:10.7554/eLife.44279.

Zhao, Y., Hull, A. K., Gupta, N. R., Goss, K. A., Alonso, J. M., Ecker, J. R., et al. (2002). Trp-dependent auxin biosynthesis in *Arabidopsis*: involvement of cytochrome P450s CYP79B2 and CYP79B3. Genes Dev. 16, 3100–3112. doi:10.1101/gad.1035402.

Zhou, N., Tootle, T. L., and Glazebrook, J. (1999). Arabidopsis *PAD3*, a gene required for camalexin biosynthesis, encodes a putative cytochrome P450 monooxygenase. Plant Cell 11, 2419–2428. doi:10.2307/3870965.

