## Supplementary Figures for "Transcriptional response to host chemical cues underpins expansion of host range in a fungal plant pathogen lineage"

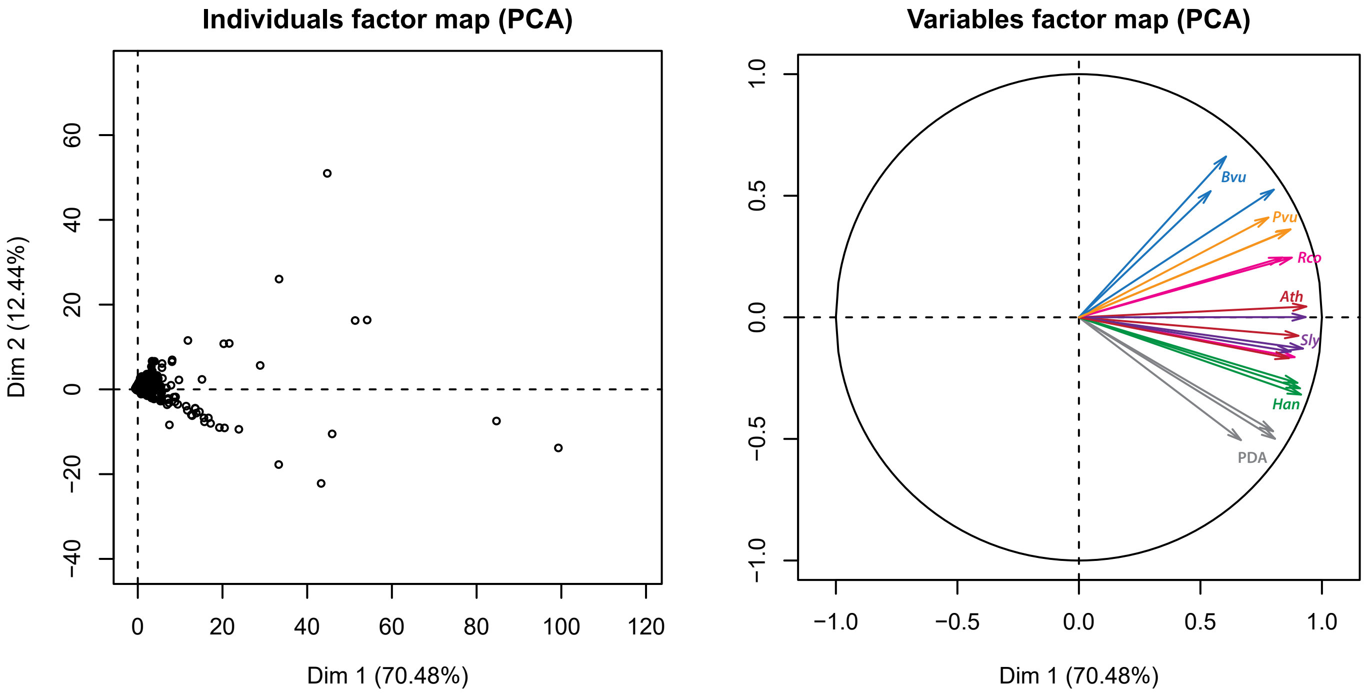


**Supplementary Figure 1. Principal component analysis of *S. sclerotiorum* gene expression during the colonization of six plants, samples harvested at the edge of invasive colonies.** PDA, Potato Dextrose Agar, *Ath, Arabidopsis thaliana; Bvu, Beta vulgaris; Han, Helianthus annuus; Pvu, Phaseolus vulgaris; Rco, Ricinus communis; Sly, Solanum lycopersicum*.

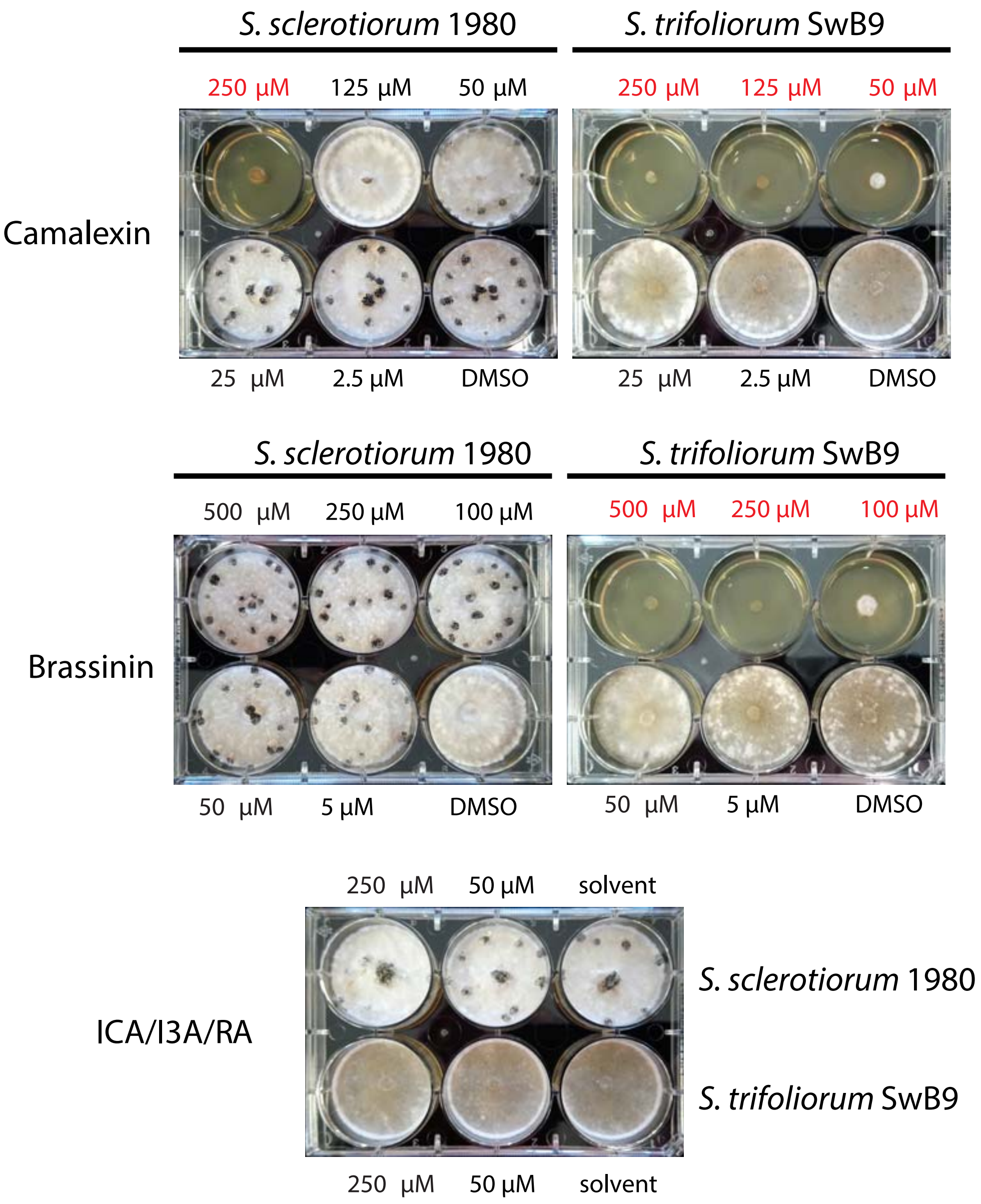


**Supplementary Figure 2. Complements to the phytoalexin *in vitro* sensitivity assay.** The assay was performed as described in Figure 5, additional concentrations and phytoalexin solutions are presented here. ICA/I3A/RA is a mixture of Indole-3 carboxylic acid (in ethanol), Indole-3 ylmethylamine (in DMSO) and raphanusamic acid (in water). Non viable concentrations are labelled in red.

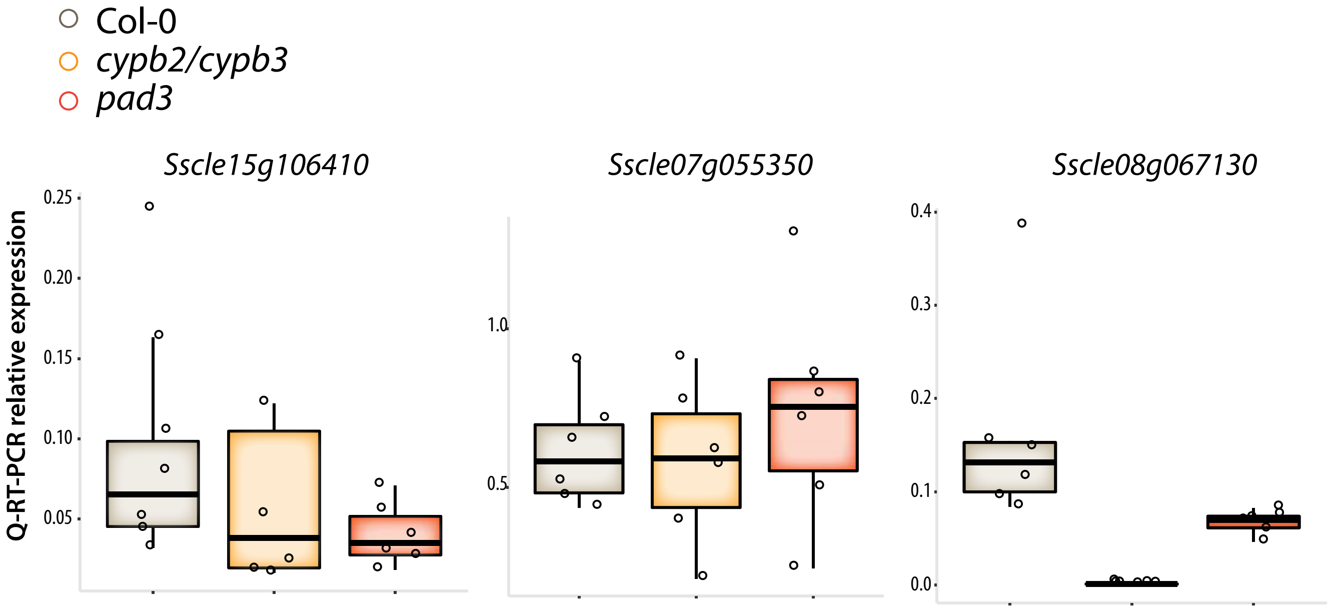


**Supplementary Figure 3. Relative expression at 72 hours post inoculation for 3 *S. sclerotiorum* genes** determined by quantitative reverse transcription PCR (Q RT-PCR) on *A. thaliana* wild type plants, *cypb2/cypb3* and *pad3* mutants. Values shown are for 6 independent biological replicates averaged over two technical replicates.

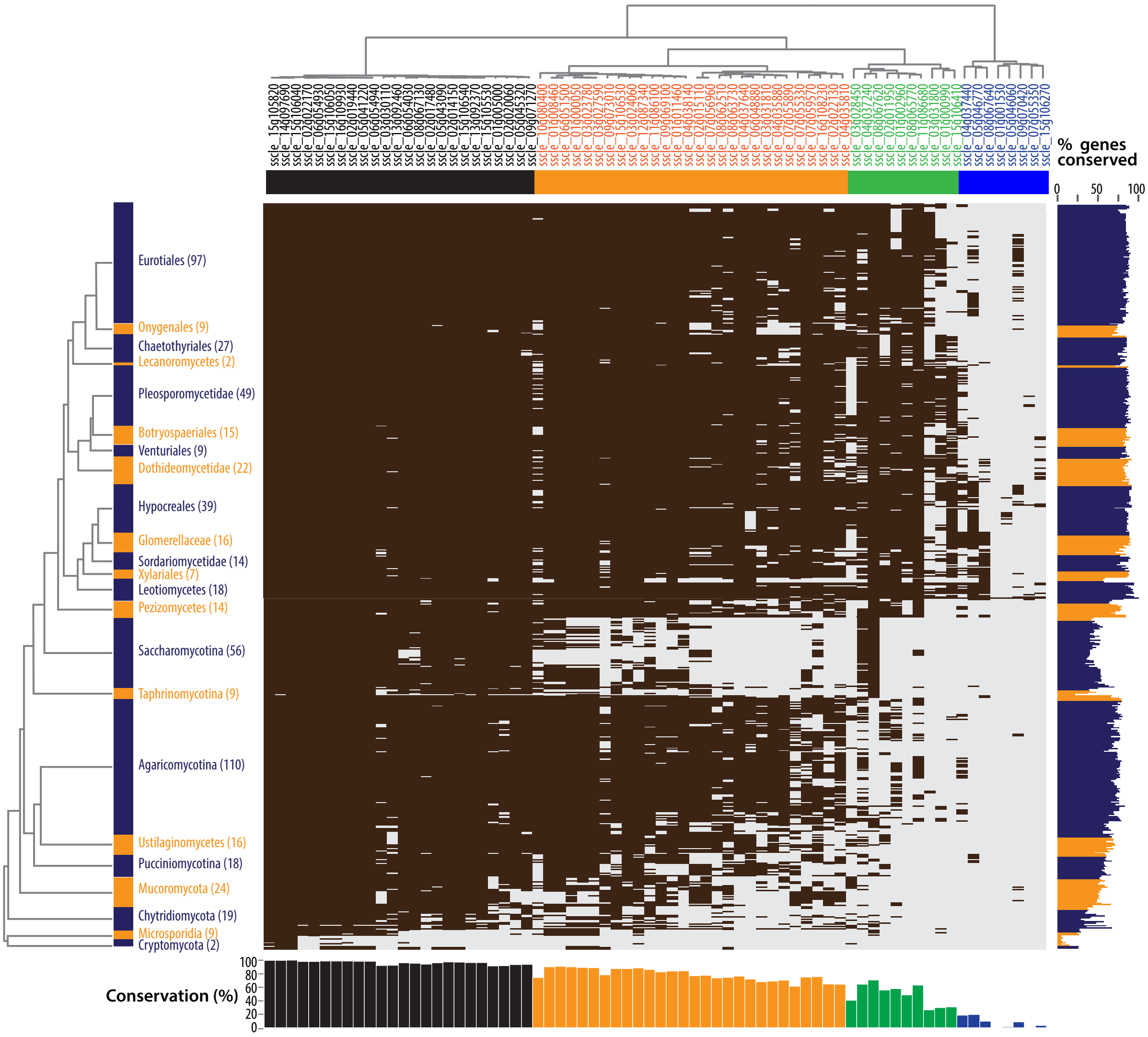


**Supplementary Figure 4. Phylogenetic distribution of the 70 *S. sclerotiorum* genes upregulated during *A. thaliana* infection and on camalexin.** Conservation of *S. sclerotiorum* genes (columns) in 629 fungal species (rows) is shows as a brown box when a BlatsP hit with p-value < 1E-10 was detected, as light grey boxes otherwise. Fungal species are organized following the species phylogeny retrieved from JGI Mycocosm, number in brackets indicate the number of species per lineage. The % of conserved genes bars show the proportion of genes detected in each species. The conservation % bars show the proportion of fungal species in which each gene was detected.

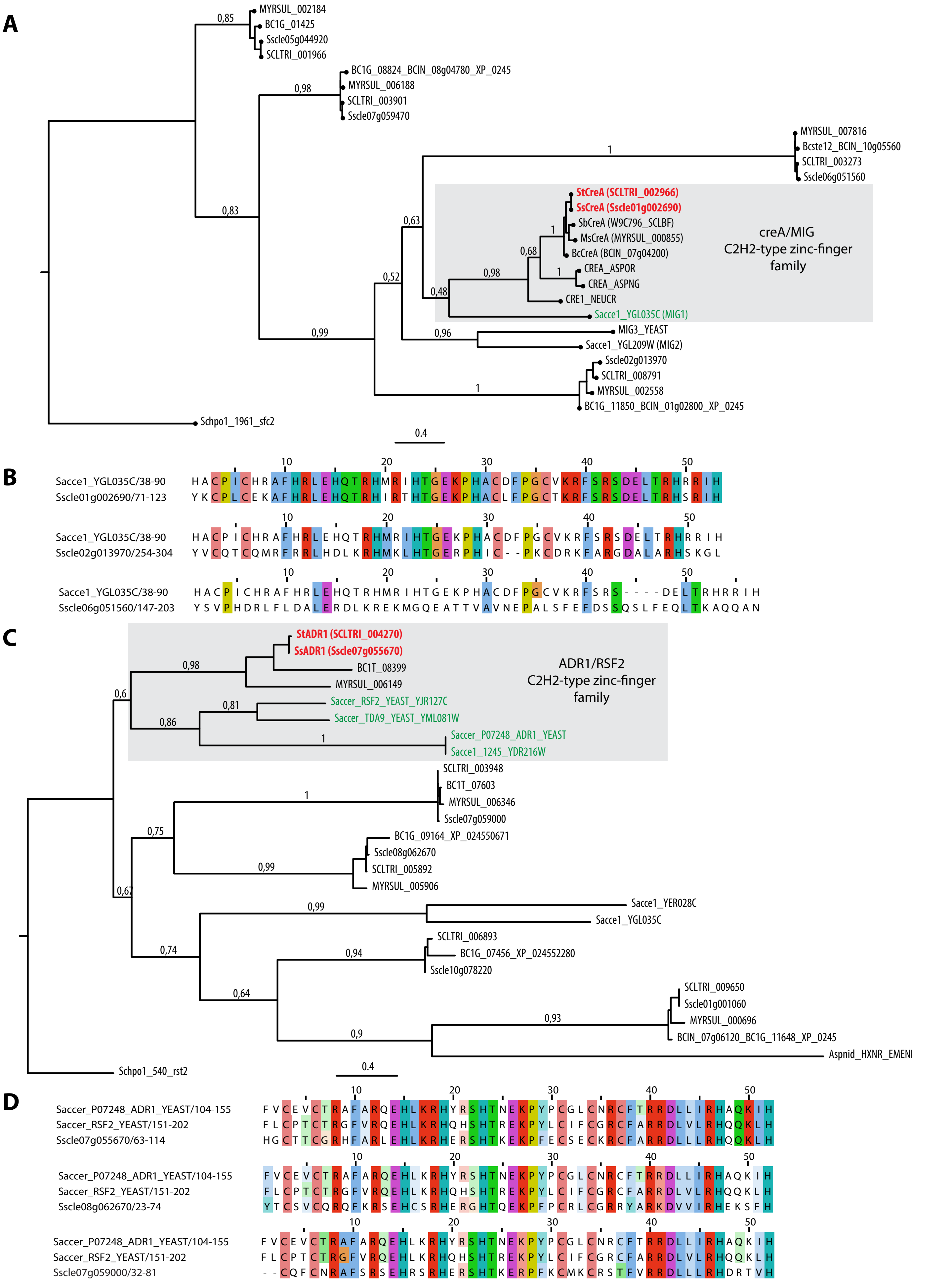


**Supplementary Figure 5. Phylogenetic analysis of *Sclerotinia* orthologs of yeast *Mig1-3* and *Adr1/Rsf2* genes. (A, C)** Maximum likelihood phylogenetic trees obtained with PhyML using the WAG model for amino acid substitutions. The trees were rooted on *Schizosaccharomyces pombe* closest homolog. Labels show branch support determined by an approximate likelihood ratio test. **(B, D)** Multiple sequence alignments of the zinc finger regions for *S. sclerotiorum* and *Saccharomyces cereviseae* homologs. Conserved residues are colored according to % conservation and residue type.

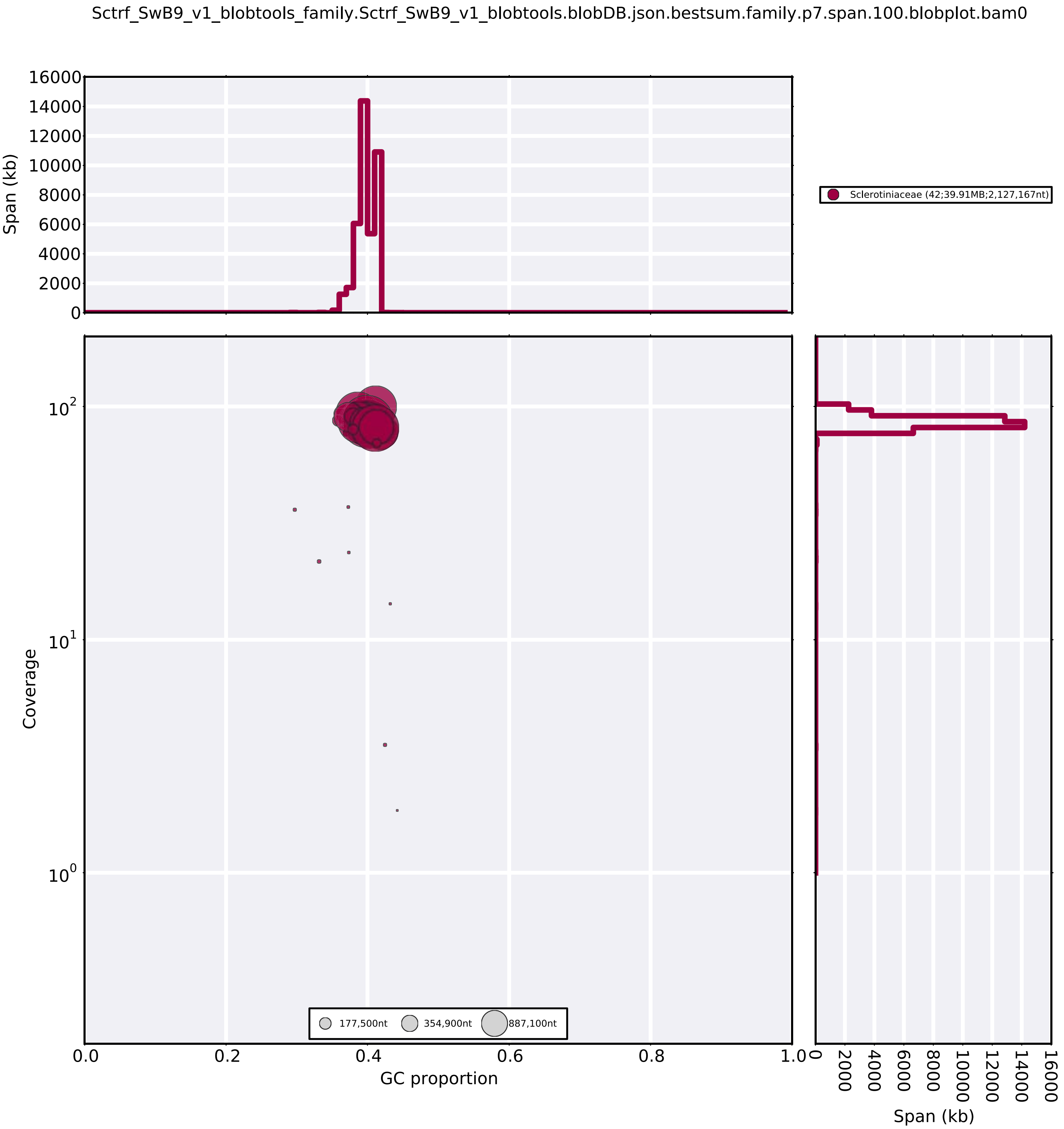


**Supplementary Figure 6. Blobtools analysis for *S. trifoliorum* genome assembly.**
